# Novel, small molecules targeting the 5-HT_4_ receptor protect against stress-induced maladaptive behavior with efficacy across age

**DOI:** 10.1101/2025.10.08.680712

**Authors:** Rebecca Ravenelle, Shi-Xian Deng, Mel Nelson, Maria H. Pedersen, Indira Mendez-David, Denis J. David, Victor M. Luna, Gergely Turi, Madeline Abramson, Jonathan A. Javitch, Donald W. Landry, Briana K. Chen, Christine A. Denny

## Abstract

**BACKGROUND:** Stress is a risk factor for developing psychiatric disorders, including major depressive disorder (MDD). Compounds targeting the serotonin type 4 receptor (5-HT_4_R) hold promise as novel rapid-acting treatments of mood disorders. However, a lack of selectivity and numerous side effects have been limiting factors for their clinical use. Here, we developed and characterized novel-composition 5-HT_4_R compounds in mouse models of stress.

**METHODS:** Three 5-HT_4_R-targeting compounds were designed and synthesized based on PF-04995274, a high-affinity 5-HT_4_R ligand reported to be a partial agonist. G-protein assays were utilized to characterize molecular activity. Saline, PF-04995274, or a novel compound were administered before or after stress in male and female mice. Drug effects were assayed using behavioral paradigms. Patch clamp electrophysiology was used to determine the effect of drug on glutamatergic activity in hippocampal Cornu Ammonis 3 (CA3).

**RESULTS:** Prophylactic administration of DL5, DL7, or DL8 was effective at reducing stress-induced maladaptive behaviors in male and female mice; DL7 and DL8 were effective when administered after stress. When administered following learned helplessness (LH), DL7 reduced behavioral despair and increased c-Fos in the dentate gyrus (DG) and CA3. All novel compounds attenuated large-amplitude AMPA receptor–mediated bursts in ventral CA3 (vCA3). In aged male mice, prophylactic DL7 reduced behavioral despair.

**CONCLUSIONS:** These results characterize novel 5-HT_4_R-targeting compounds for stress-induced psychiatric disease with the potential to address unmet needs in adult and aged patients with stress-induced psychiatric illness. Future work will characterize their mechanism of action with the goal of clinical development.

## INTRODUCTION

Depressive disorders impact over 65 million people worldwide making them a leading cause of disability (1,2). Those diagnosed with major depressive disorder (MDD) exhibit high comorbidity with other psychiatric disorders, such as generalized anxiety disorder (GAD) (3), which is associated with reduced symptom remission following pharmacological treatment (4). Additionally, there is a health disparity in gender; women are twice as likely as men to be diagnosed with MDD or GAD during their lifetime (5–7). Stressful or traumatic life events are a major risk factor for symptom onset and severity (8–11).

In adults with MDD, only half receive treatment (12). Many individuals report worsening of symptoms upon treatment onset (13–16) and about 31% of patients do not achieve remission after full courses of two or more antidepressants (17) indicating a need for new pharmacotherapies. Recent work from our group and others has identified the serotonin type 4 receptor (5-HT_4_R) as a promising target to treat depression and anxiety (18–22). The 5-HT_4_R is a G protein-coupled receptor that activates the cyclic adenosine monophosphate (cAMP) – protein kinase A (PKA) pathway, resulting in excitation of neurons in response to serotonin. 5-HT_4_Rs are found throughout regions implicated in anxiety and depression, including the hippocampus (HPC), medial prefrontal cortex, amygdala (AMG), and striatum (23–26).

In previous preclinical work, subchronic administration of the 5-HT_4_R partial agonist RS-67,333 reduced anxiety-like behavior and behavioral despair following corticosterone (CORT) exposure (27), and acute administration of RS-67,333 or the 5-HT_4_R agonist prucalopride also decreased behavioral despair, (28) indicating 5-HT_4_R activation produces rapid-acting antidepressant-like effects. Additionally, our group recently identified a role for 5-HT_4_R ligands in enhancing stress resilience. A prophylactic injection of prucalopride or PF-04995274 attenuated learned fear and decreased behavioral despair in male mice. Moreover, prophylactic administration of prucalopride reduced large AMPA receptor-mediated bursts in CA3, highlighting a common mechanism with the fast-acting antidepressant and prophylactic (*R,S*)-ketamine (18,29).

Treatment options selectively targeting the 5-HT_4_R are limited. Expression of 5-HT_4_Rs occurs throughout the gastrointestinal (30–32) and cardiovascular system (33,34), making peripheral side effects common. However, there is an effort to identify 5-HT_4_R compounds that will improve psychiatric symptoms. In the United States, prucalopride is approved for chronic idiopathic constipation (35,36) and recent work in healthy volunteers found that subchronic administration improves memory and alters neural activity, particularly in the HPC (37,38). Recently, a clinical trial was conducted to determine if the purported partial agonist PF-04995274 improves emotional processing and neural activity in individuals with treatment-resistant depression (TRD) (NCT03515733) (39).

As there are currently no FDA-approved 5-HT_4_R agonists to treat psychiatric symptoms, here, we sought to develop and characterize novel, small molecules targeting the 5-HT_4_R. Compounds were characterized using cAMP inhibition and G protein recruitment assays. To determine if these novel compounds were prophylactic against stress, we administered the compounds – DL5, DL7, or DL8 – to male and female mice prior to acute stress. As stressful experiences are often unpredictable, necessitating post-stress treatment, we also administered compounds *following* stress. Slice electrophysiology and immunohistochemistry against the immediate early gene (IEG) c-Fos (40) were used to explore how these compounds influenced HPC activity. Overall, by utilizing a diverse set of behavioral paradigms, we identified novel compounds that improve stress-induced maladaptive behaviors. Altogether, our data suggest these novel drugs show promise as future treatments for psychiatric disorders.

## METHODS AND MATERIALS

For a full description of Methods and Materials, please refer to the **Supplemental Methods** in Supplement 1.

### Mice

Male and female 129S6/SvEvTac mice were purchased from Taconic (Hudson, NY) at 7 – 8 weeks of age or bred in house. Male and female C57BL/6J were bred in house. Males and females were utilized for behavioral experiments and were conducted with adult (9-week-old) or aged (17-month-old) mice. Electrophysiological experiments utilized adult male 129S6/SvEvTac mice. Mice were housed 4-5/cage in a 12-h (06:00 – 18:00) light-dark colony room at 22°C with food and water available *ad libitum*. Behavioral tests were conducted during the light phase. All procedures were conducted in accordance with the National Institutes of Health (NIH) regulations and approved by the Institutional Animal Care and Use Committees (IACUCs) of the New York State Psychiatric Institute (NYSPI) and Columbia University Irving Medical Center (CUIMC).

## RESULTS

### Synthesis of novel 5-HT_4_R-targeting compounds

Based on the structure of the purported 5-HT_4_R partial agonist, PF-04995274, we designed and synthesized novel chemical entities DL5, DL7, and DL8 (**D**onald **L**andry) likely to modulate the 5-HT_4_R.

### Novel compounds exhibit variable affinity for 5-HT_4_Rs and function as weak or inverse agonists

The 5-HT_4_R is reported to have constitutive activation of G_s_ and stimulation of adenylate cyclase (41,42). To measure potency and efficacy of our novel ligands, we created a 5-HT_4_R fused at its C-terminus with Nanoluc and set up a direct recruitment assay with miniGs fused to mVenus. miniGs is a G_s_ protein engineered to reside in the cytoplasm until recruited to an activated receptor at the membrane and measures a direct interaction between receptor and activated G protein (**Figure 1A**) (43). This led to highly reproducible results, with prucalopride behaving as a partial agonist with E_max_ ∼80% of 5-HT, and RS-67,333 behaving as a more potent partial agonist with E_max_ ∼25% of 5-HT. PF-04995274, which is purportedly a partial agonist (44), behaved as an inverse agonist, lowering the E_max_ to ∼20% below vehicle baseline (**Figure 1B**). Novel compounds DL5, DL7, and DL8 were without apparent effect (**Figure 1B**) making it difficult to determine whether compounds were acting at the receptor as neutral antagonists or were inactive.

**Figure 1.**
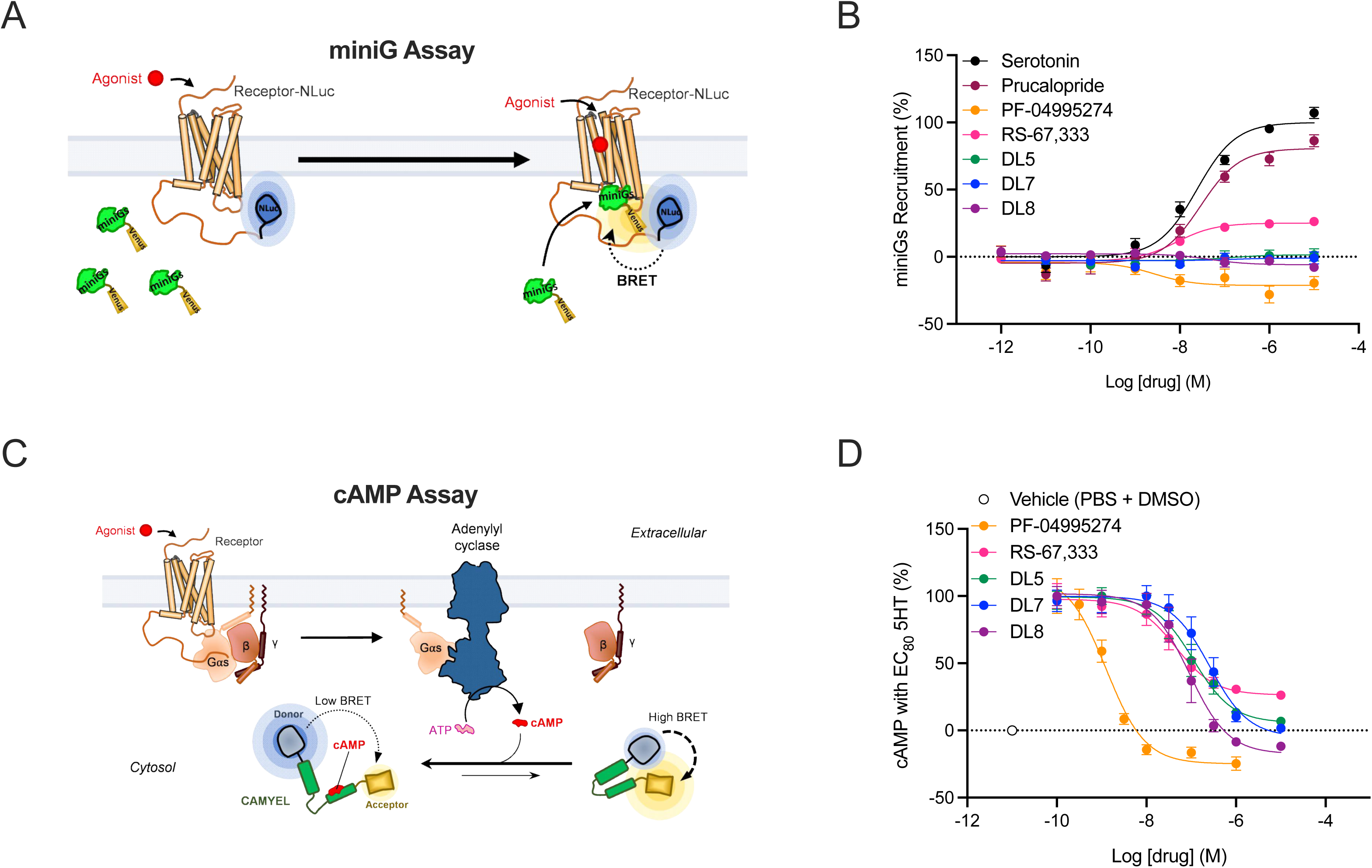
Activity of novel 5-HT_4_R-targeting compounds in direct miniG and a cAMP accumulation competition assay. **(A)** Schematic of the miniGs direct recruitment assay, showing the 5-HT_4_R fused to nanoluciferase at its C-terminus and miniGs fused to the acceptor mVenus. Receptor activation due to ligand binding results in recruitment of miniG proteins from the cytoplasm to the receptor at the plasma membrane leading to an increase in BRET between the donor and acceptor molecules. **(B)** Dose response curves for the miniGs assays with standards serotonin, prucalopride, partial agonist RS-67,333, and parental compound PF-04995274, which acts as an inverse agonist. Compounds DL5, DL7, and DL8 have minimal effects in this system. **(C)** Schematic of the cAMP assay, showing activation of the Gα subunit of the G_s_ heterotrimeric complex in response to receptor activation, resulting in an increase in intracellular cAMP. Changes in intracellular cAMP are monitored using the cAMP sensor CAMYEL, an EPAC protein that is fused to both a donor luciferase and acceptor fluorescent protein. As cAMP is produced, it binds to EPAC and causes conformational changes that lead to a decrease in the BRET signal. The experiments in **(D**) are competition assays against an EC80 concentration of serotonin (35 nM). RS-67,333 decreases residual cAMP levels in a concentration dependent manner, replacing serotonin but acting as a partial agonist. PF-04995274 also leads to concentration dependent decrease in cAMP levels but to a value below baseline, consistent with its actions as an inverse agonist. Compounds DL5 and DL7 lead to concentration dependent decreases in residual cAMP to near baseline values. Compound DL8 lead to decreases in residual cAMP below the vehicle baseline similar to PF-04995274. Data represented as a percentage normalized to E_max_ of serotonin (**D**) or EC80 serotonin (**D**). Dose response curves were fit using a 3-parameter model and values represent the mean of 3 – 7 individual replicates. Each replicate was performed with triplicate determinations. Error bars represent ± SEM; cAMP, cyclic adenosine monophosphate; EC80, effective concentration; 80% nluc, nanoluciferase.

To better elucidate the activity of these compounds, we conducted an inhibition mode assay, competing with an EC80 concentration of the full agonist 5-HT (**Figure 1C**). We obtained reliable inhibition curves for RS-67,333 that plateaued at about 25%, consistent with its action as a partial agonist (**Figure 1D**). PF-04995274 again behaved as an inverse agonist. Compounds DL5 and DL7 inhibited completely, consistent with neutral antagonism, whereas DL8 behaved as an inverse agonist like the parental compound (**Figure 1D**). These data confirm that these novel compounds interact with the 5-HT_4_R and alter downstream activity, which may ultimately influence neural circuits underlying behavior.

### A prophylactic injection of novel 5-HT_4_R compound DL5, DL7 or DL8 protects against stress

Previously, we have shown that a single prophylactic administration of prucalopride or PF-04995274 attenuates learned fear and behavioral despair in male mice (18). Here, we sought to determine if the novel 5-HT_4_R compounds are also effective at preventing maladaptive behaviors. We utilized 129/SvEvTac mice as this strain exhibits greater susceptibility to stress (46,48). Adult male 129S6/SvEvTac mice were administered saline, PF-04995274 (10 mg/kg), or DL5, DL7, or DL8 (3 or 10 mg/kg) (**Figure 2A**). Dosing was based on prior studies of 5-HT_4_R ligands (18). One week later, mice were administered 3-shock contextual fear conditioning (CFC) as a stressor. Drug administration did not alter freezing during CFC training (**Figure 2B-C**). Five days later, mice were returned to the context for a retrieval session. Prophylactic PF-04995274, as previously demonstrated (18), and DL7 (3 mg/kg) attenuated learned fear (**Figure 2D-E**).

**Figure 2.**
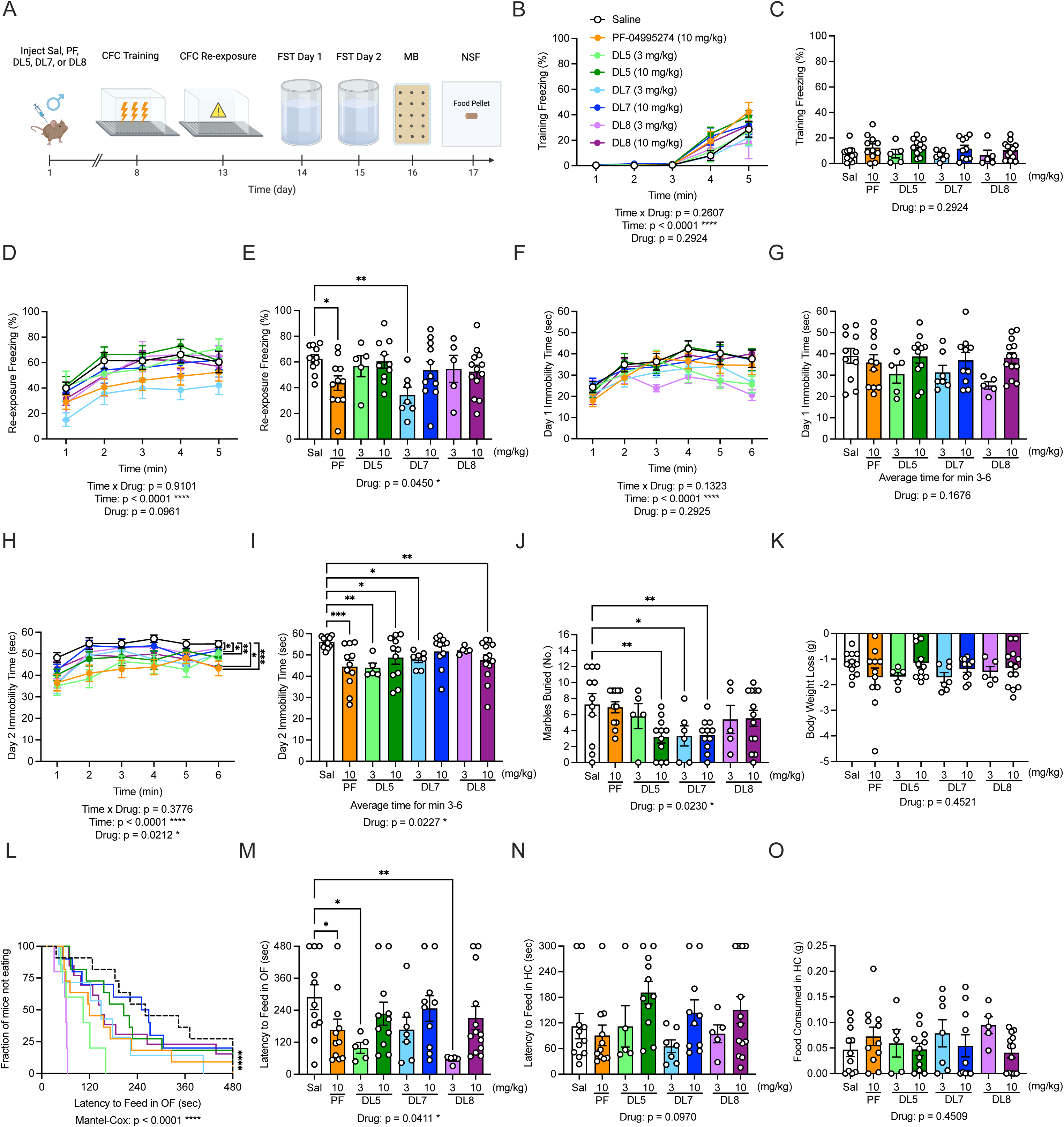
A single prophylactic injection of a novel 5-HT_4_R compound protects against stress-induced maladaptive behavior in male mice. **(A)** Experimental protocol. **(B-C)** Freezing was comparable across all groups during CFC training **(B)** across time and **(C)** averaged across the entire test session. **(D-E)** During context re-exposure, PF-04995274 (10 mg/kg) and DL7 (3 mg/kg) decreased fear expression when compared with saline mice. **(F-G)** On day 1 of the FST, immobility time was comparable across groups. **(H-I)** On day 2 of the FST, PF-04995274, DL5 (3 and 10 mg/kg), DL7 (3 mg/kg), and DL8 (10 mg/kg) decreased immobility time when compared with saline. **(J)** In the MB task, DL5 (10 mg/kg) and DL7 (3 and 10 mg/kg) decreased the number of marbles buried when compared to saline. **(K)** Prior to administration of NSF, mice lost a comparable amount of weight during food restriction across groups. **(L-M)** In the NSF, PF-04995274 (10 mg/kg), DL5 (3 mg/kg), and DL8 (3 mg/kg) reduced the latency to feed in the OF. In the home cage, groups exhibited comparable **(N)** latency to feed and **(O)** amount of food consumed. Sal n = 11, PF n = 11, DL5 (3 mg/kg) n = 5, DL5 (10 mg/kg) n = 11, DL7 (3 mg/kg) n = 7, DL7 (10 mg/kg) n = 10, DL8 (3 mg/kg) n = 5, DL8 (10 mg/kg) n = 13, male mice per group; error bars represent ± SEM; *p < 0.05, ** p < 0.01, *** p < 0.001. CFC, contextual fear conditioning; FST, forced swim test; HC, home cage; MB, marble burying; NSF, novelty-suppressed feeding; OF, open field; PF, PF-04995274; Sal, saline. Behavioral timeline created with BioRender.com.

On FST day 2, but not day 1, PF-04995274 reduced immobility time, consistent with previous findings (18). DL5 (3 and 10 mg/kg), DL7 (3 mg/kg), and DL8 (10 mg/kg) also reduced immobility time (**Figure 2F-I**). In the marble burying (MB) task, DL5 and DL7 reduced the number of marbles buried, indicating a reduction in perseverative behavior (**Figure 2J**). In the novelty suppressed feeding (NSF) task, PF-04995274, DL5, and DL8 reduced the latency to feed in the open field (OF) arena (**Figure 2K-O**). To determine if prophylactic administration impacts behavior in the absence of a stressor, we prophylactically administered PF-04995274 (10 mg/kg), DL5, DL7, or DL8 (3 mg/kg) (Figure S1A) and then administered no-shock CFC, MB, open field (OF), and NSF. Prophylactic drug administration did not impact behavior compared to saline (Figure S1B-O). Together these data highlight the prophylactic efficacy of DL5, DL7, or DL8 to prevent stress-induced maladaptive behaviors.

Adult female 129S6/SvEvTac mice were administered saline, PF-04995274 (3 mg/kg), or DL5, DL7, or DL8 (3 mg/kg) (**Figure S2A**). Again, dosing of PF-04995274 was based on prior studies (18). Drug administration did not alter freezing during CFC training or context re-exposure (**Figure S2B-E**). On FST day 1, DL8 increased immobility time in female mice (**Figure S2F-G**). On FST day 2, all mice exhibited comparable immobility time (**Figure S2H-I**). In the MB task, DL7 and DL8 significantly reduced the number of marbles buried (**Figure S2J**). In the NSF task, PF-04995274 and DL5 reduced the latency to feed in the OF (**Figure 2K-O**). These data demonstrate the prophylactic efficacy of DL7 and DL8 for perseverative behavior and DL5 for hyponeophagia in female mice.

### Novel 5-HT_4_R compounds DL7 and DL8 are effective when administered following stress in male mice

We then sought to determine efficacy of these compounds when administered *following* acute stress (**Figure 3A**). Five minutes following 3-shock CFC, 129S6/SvEvTac mice were administered saline, PF-04995274 (10 mg/kg), or DL5, DL7, or DL8 (3 or 10 mg/kg). All groups froze comparably during training (data not shown), and drug administration did not alter freezing during context re-exposure (**Figure 3B-C**). During FST day 1, DL5, DL7, and DL8 increased immobility time (**Figure 3D-E**). During FST day 2, PF-04995274, DL7 (10 mg/kg), and DL8 (10 mg/kg) decreased immobility time (**Figure 3F-G**). Drug administration did not alter behavior in the MB (**Figure 3H**) or NSF tasks (**Figure 3I-3M**).

**Figure 3.**
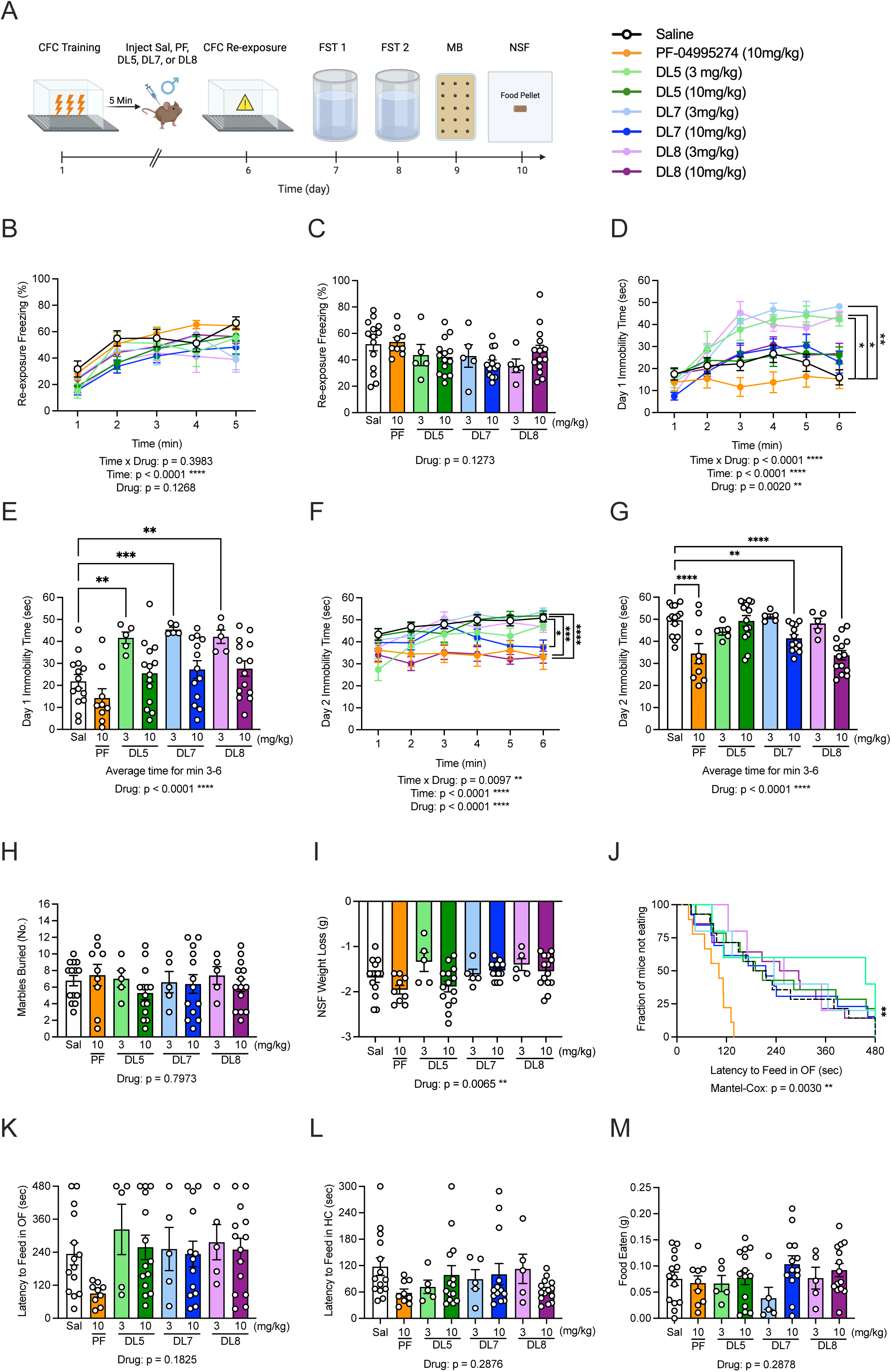
A single injection of a novel 5-HT_4_R compound decreases behavioral despair when administered following stress in male mice. **(A)** Experimental design. **(B-C)** Freezing was comparable across all groups during CFC exposure. **(D-E)** On day 1 of the FST, DL5, DL7, and DL8 (all 3 mg/kg) increased immobility time. **(F-G)** On day 2 of the FST, PF-04995274 (10 mg/kg), DL7 (10 mg/kg), and DL8 (10 mg/kg) reduced immobility time when compared with saline. **(H)** In the MB task, all groups buried a comparable number of marbles. **(I)** In the NSF, all groups lost a similar amount of weight when compared directly to saline although there was an overall effect of Drug. **(J-K)** During the NSF, there was an overall effect of Drug, but no drug decreased the latency to feed in the OF when compared to saline. In the home cage, **(L)** latency to feed and **(M)** food consumed was comparable across all groups. Sal n = 14, PF n = 9, DL5 (3 mg/kg) n = 5, DL5 (10 mg/kg) n = 14, DL7 (3 mg/kg) n = 5, DL7 (10 mg/kg) n = 13, DL8 (3 mg/kg) n = 5, DL8 (10 mg/kg) n = 14 male mice per group; error bars represent ± SEM; * p < 0.05, ** p < 0.01, *** p < 0.001. CFC, contextual fear conditioning; FST, forced swim test; MB, marble burying; NSF, novelty suppressed feeding; OF, open field; HC, home cage; Sal, saline; PF, PF-04995274; min, minutes; sec, seconds; mg, milligram; kg, kilogram. Behavioral timeline created with BioRender.com.

In female 129S6/SvEvTac mice, five minutes following 3-shock CFC, mice were administered saline, PF-04995274 (10 mg/kg), or DL5, DL7, or DL8 (3 or 10 mg/kg) (**Figure S3A**). All groups froze comparably during training (data not shown), and drug administration did not alter freezing during context re-exposure (**Figure S3B-C**). During the FST and MB, all mice exhibited comparable behavior (**Figure S3D-H**). In the NSF, PF-04995274, DL5, and DL8 decreased the latency to feed in the OF arena (**Figure S3I-M**). Together our data indicate these compounds are efficacious for behavioral despair in males and hyponeophagia in females when administered shortly following a stressor.

### 5-HT4R compounds decrease perseverative behavior in C57BL/6J female, but not male mice

To extend these findings to other mouse strains, we administered PF-04995274 or prucalopride (3 or 10 mg/kg) following stress in C57BL/6J mice, a more resilient strain (45,47,49). Neither drug improved stress-induced maladaptive behavior in male mice; unexpectedly PF-04995274 increased perseverative behavior in the MB task (**Figure S4**). In C57BL/6J female mice, PF-04995274 (3 mg/kg) and prucalopride (10 mg/kg) decreased the number of marbles buried (**Figure S5**). Here, acute administration of a 5-HT_4_R agonist reduces perseverative behavior in females only, in a mouse line resilient to stress.

### Novel 5-HT_4_R compounds alter AMPA receptor-driven synaptic bursts in CA3

We have previously shown that prophylactic administration of (*R,S*)-ketamine, (2*R*,6*R*)-hydroxynorketamine (HNK), prucalopride, and fluoroethylnormemantine (FENM) reduce large AMPA receptor (AMPAR)-mediated excitatory postsynaptic currents (EPSCs) in CA3 (18,50,51) To determine if the new compounds share this effect, we administered saline, DL5, DL7, or DL8 to adult male 129S6/SvEvTac mice. One week later, mice were euthanized, and we conducted whole-cell voltage clamp recordings of spontaneous EPSCs in CA3 (**Figure 4A**). DL5, DL7, and DL8 reduced mean EPSC amplitude compared to saline (-46.34 ± 1.33 pA) (**Figure 4B-E, F**).

**Figure 4.**
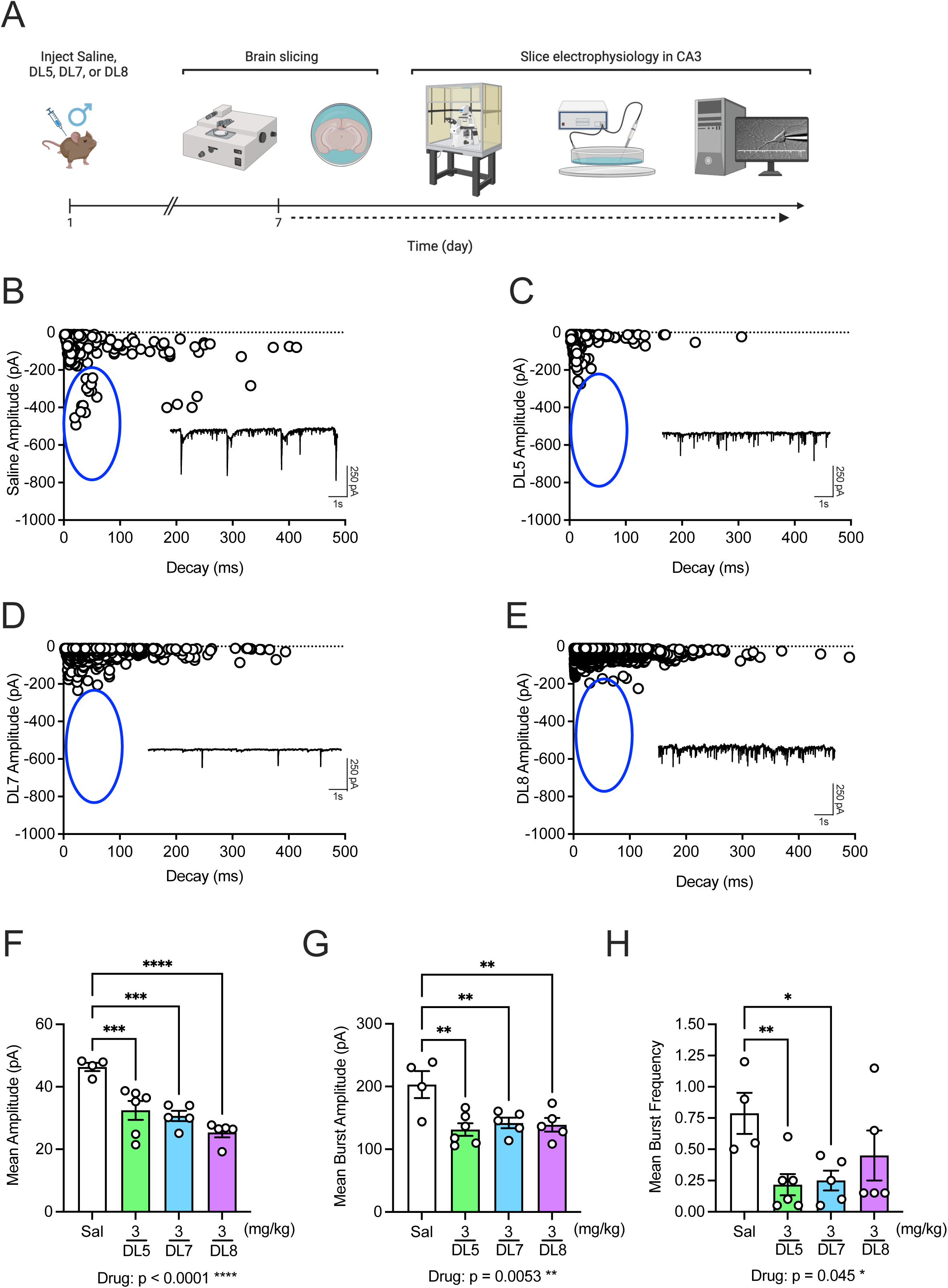
Prophylactic administration of novel 5-HT_4_R compounds reduce large AMPA-driven synaptic bursts in CA3. **(A)** Experimental design. Mice were administered saline, DL5 (3 mg/kg), DL7 (3 mg/kg), or DL8 (3 mg/kg) 1 week before whole-cell voltage clamp electrophysiology. Representative EPSCs following administration of **(B)** saline, **(C)** DL5, **(D)** DL7, and **(E)** DL8. **(F)** The mean amplitude was attenuated in DL5, DL7, and DL8-administered mice when compared to saline **(G)** The mean burst amplitude was significantly reduced by DL5, DL7, and DL8 when compared with saline. **(H)** The mean frequency of all AMPAR-mediated EPSCs within a 20-second recording period was significantly decreased by DL5 and DL7, but not by DL8, when compared with saline. Sal n = 4, DL5 n = 6, DL7 n = 5, DL8 n = 5 cells per group; error bars represent ± SEM; *p < 0.05, ** p < 0.01, *** p < 0.001, **** p < 0.0001. AMPAR, AMPA receptor; CA3, cornu ammonis area 3; EPSC, excitatory postsynaptic current; Sal, saline; ms, millisecond; pA, picoampere. Behavioral timeline created with BioRender.com.

Novel compounds also reduced mean amplitude of bursts compared to controls (203.20 ± 21.48) (**Figure 4G**) while DL5 and DL7 also reduced mean burst frequency (**Figure 4H**). These data indicate that in addition to reducing behavioral despair, prophylactic administration of novel compounds may modify AMPA-R signaling to enhance resilience.

### Novel compound DL7 is antidepressant when administered following learned helplessness

To extend the capacity of these compounds for other stressors, we next utilized LH (52–55). As prior experiments failed to identify improvement in behavioral despair in females, only male 129S6/SvEvTac mice were utilized. Five minutes following LH training (**Figure 5A**), mice were administered saline, PF-04995274, or DL5, DL7, or DL8 (10 mg/kg). Five days later, mice underwent a shock escape protocol. There were no group differences in activity during the habituation period, total session latency, or latency to escape (**Figure 5B–E**).

**Figure 5.**
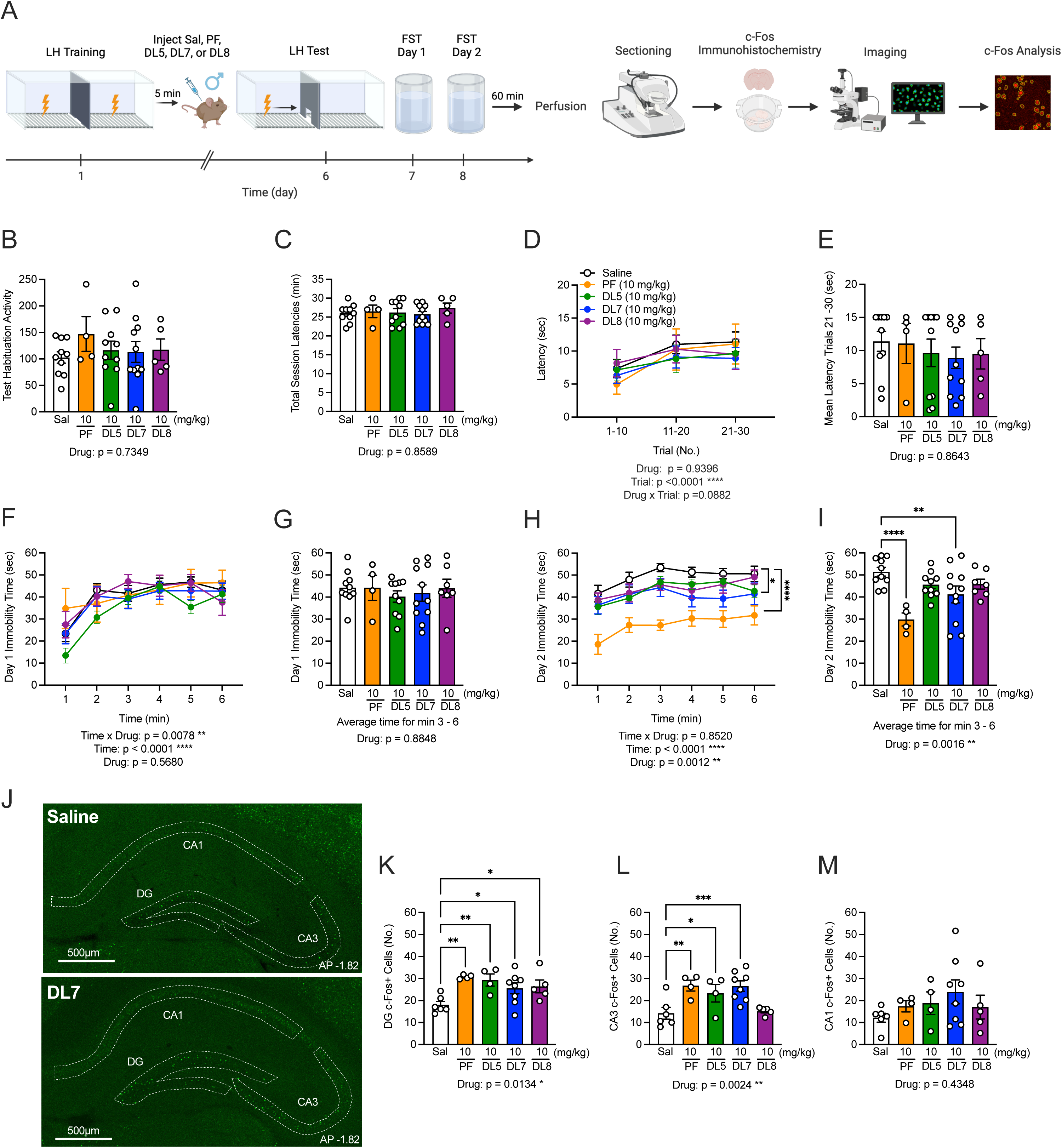
DL7 reduces behavioral despair when administered after learned helplessness in male mice. **(A)** Experimental design. **(B)** Total activity during habituation period was comparable across all groups. **(C)** Total session latencies were comparable across all groups. **(D)** Mean latency to escape the shock was comparable across all groups. **(E)** Mean latency to escape in trials 21 – 30 was comparable across all groups. **(F-G)** On day 1 of the FST, all groups exhibited comparable immobility time compared to saline. **(H-I)** On day 2 of the FST, PF-04995274 and DL7 reduced immobility time compared with saline. **(J)** Representative images of c-Fos expression (green) in the HPC; scale bar = 500 µm. **(K)** PF-04995274, DL5, DL7, and DL8 increased c-Fos expression in the DG when compared with saline. **(L)** PF-04995274, DL5, and DL7 increased c-Fos expression in CA3 when compared with saline. **(M)** c-Fos expression was comparable across groups in CA1. Sal n = 10, PF n = 4, DL5 n = 10, DL7 n = 11, DL8 n = 5 male mice per group; error bars represent SEM; * p < 0.05, ** p < 0.01, *** p < 0.001. CFC, contextual fear conditioning; ; CA3, cornu ammonis area 3; CA1, cornu ammonis area 1; DG, dentate gyrus; FST, forced swim test; HC, home cage; HPC, hippocampus; MB, marble burying; NSF, novelty suppressed feeding; OF, open field; PF, PF-04995274; Sal, saline; min, minutes; sec, seconds; mg, milligram; kg, kilogram. Behavioral timeline created with BioRender.com.

On FST day 1, all groups exhibited comparable immobility time (**Figure 5F-G**). On FST day 2, PF-04995274 and DL7 reduced immobility time (**Figure 5H-I**). Sixty minutes following the FST, mice were euthanized, and brains were processed for neural activity using an antibody against c-Fos (**Figure 5J**). Administration of all drugs increased c-Fos expression in the DG (**Figure 5K**). PF-04995274, DL5, and DL7 increased c-Fos expression in CA3 (**Figure 5L**). Interestingly, DL8 did not increase c-Fos expression in CA3 and this compound also failed to reduce mean EPSC amplitude in CA3 in contrast to other novel compounds. Drug administration did not alter c-Fos expression in CA1 (**Figure 5M**). Together, this data suggests these drugs produce long-lasting neural changes in the DG and CA3, regions we have previously shown are modulated by prophylactic (*R,S*)-ketamine (51,56).

### Prophylactic DL7 attenuates stress-induced behavioral despair in aged male mice

We previously found that the (*R,S*)-ketamine is not effective at improving stress-induced behavioral despair in aged mice (57,58). This parallels clinical findings that antidepressant treatments lack efficacy in older adults (59–62). To determine if these compounds are effective in aged 129S6/SvEvTac mice, we administered RS-67,333, prucalopride, PF-04995274, or DL7 (3 or 10 mg/kg) one week prior to CFC (**Figure 6A**). In male mice, drug administration did not influence freezing behavior during CFC training or retrieval (**Figure 6B-E**). On FST day 1, all groups exhibited comparable immobility (**Figure 6F-G**). One FST day 2, RS-67,333, PF-04995274, and DL7 reduced immobility time (**Figure 6H-I**). There was no effect of drug administration in the MB or NSF assays (**Figure 6J-O**).

**Figure 6.**
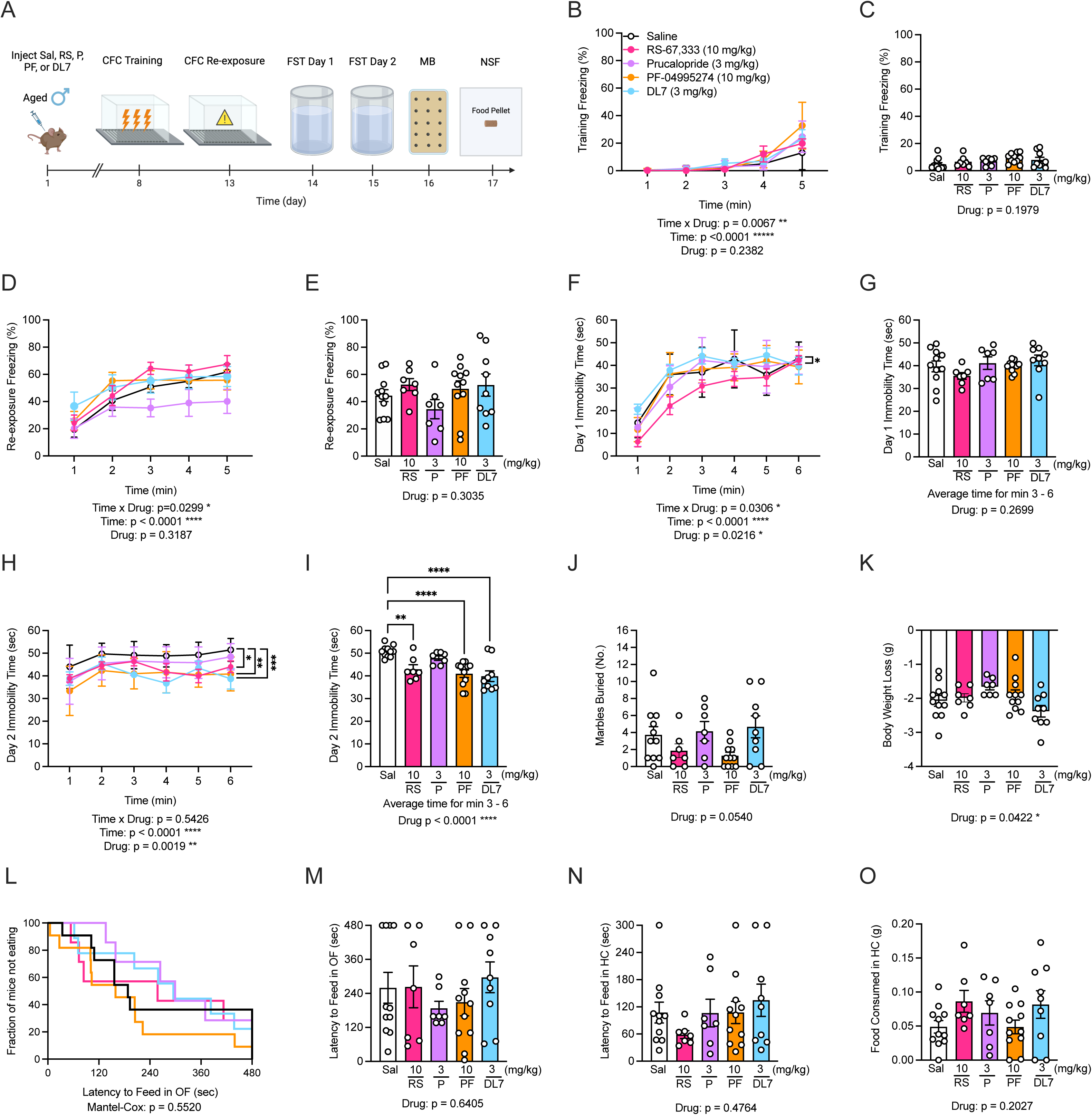
Prophylactic DL7 exerts therapeutic potential in aged male mice. **(A)** Experimental timeline for aged (∼17 month) male mice. **(B-C)** Freezing during CFC training was comparable across groups. **(D-E)** During CFC exposure, freezing behavior was comparable across groups. **(F)** On day 1 of the FST, RS-67,333 reduced immobility across the entire experimental session compared to saline, but **(G)** average immobility time for 3 – 6 min was comparable across groups. **(H-I)** On day 2 of the FST, RS-67,333 (10 mg/kg), PF-04995274 (10 mg/kg), and DL7 (3 mg/kg) reduced immobility compared to saline. **(J)** In the MB task, all groups buried a similar number of marbles compared to saline. **(K)** During food restriction phase of the NSF assay, drug groups lost a comparable amount of weight when compared directly to saline although there was a main effect of Drug. **(L-M)** Latency to feed in the OF in the NSF was comparable across all groups. In the home cage, **(N)** latency to feed and **(O)** the amount of food consumed was comparable between the groups. Sal n = 11, RS n = 7, Pruc n = 7, PF n = 11, DL7 n = 9, male mice per group; error bars represent ± SEM; * p < 0.05, ** p < 0.01, *** p < 0.001. CFC, contextual fear conditioning; FST, forced swim test; MB, marble burying; NSF, novelty suppressed feeding; OF, open field; HC, home cage; Sal, saline; PF, PF-04995274; P, prucalopride; RS, RS-67,333; min, minutes; sec, seconds; mg, milligram; kg, kilogram. Behavioral timeline created with BioRender.com.

In female mice, there was no effect of drug administration on freezing during CFC training or retrieval or during FST (**Figure S6A-I**). In the MB assay, RS-67,333 (3 mg/kg), prucalopride, and PF-04995274 decreased the number of marbles buried (**Figure S6J**). In the NSF task, drug administration did not alter behavior in the OF, but RS-67,333 increased the latency to feed in the home cage (**Figure S6K-O**). Together these experiments illustrate efficacy of RS-67,333, PF-04995274, and DL7 to improve stress-induced behavioral despair in aged male mice.

## DISCUSSION

We developed novel small molecule compounds targeting the 5-HT_4_R based on the compound PF-04995274 for application to stress-induced psychiatric disease. We report that PF-04995274 behaved as an inverse agonist despite previously being reported to be a partial agonist (63,64). Results of a competing EC80 serotonin inhibition assay established that novel compounds DL5 and DL7 function as neutral antagonists, whereas DL8 functions as an inverse agonist. Despite lower affinities for the 5-HT_4_R compared to PF-04995274 *in vitro*, behavioral testing found that in male mice, prophylactic DL5, DL7, and DL8 improved stress-induced behavioral despair at similar dosing; when administered *following* stress, DL7 and DL8 were also effective. In female mice, prophylactic DL5 improved hyponeophagia while DL7 and DL8 reduced perseverative behavior; after stress, DL5 and DL8 were again effective at improving hyponeophagia. Efficacy in females at lower doses compared to males is consistent with our previous work showing sex-specific dosing of (*R,S*)-ketamine (50). A single prophylactic administration of DL5, DL7, or DL8, reduced spontaneous EPSCs in CA3, illustrating a potential mechanism by which these drugs may attenuate stress-induced maladaptive behavior. Notably, prophylactic administration of DL7 was also effective at reducing behavioral despair in aged male mice. Together this work highlights efficacy of novel 5-HT_4_R targeting compounds to prevent stress-induced maladaptive behaviors across sex and age.

We find that drug efficacy varies with administration time. For example, in adult male mice, prophylactic administration DL5 (3 and 10 mg/kg) reduces behavioral despair in FST but was ineffective if administered following CFC stress. We previously identified changes in drug efficacy with different administration time points – prophylactic administration of (*R,S*)-ketamine prior to social defeat stress improves immobility in FST, however administration of the same dose following social defeat is ineffective (68). Interestingly, when administered post-stress, several compounds increase immobility time during FST day 1. Although we determine efficacy based on day 2 of the task, in line with Porsolt *et al.,* (70), it is possible that novel compounds influence the encoding of FST on day 1. Follow-up experiments may shed light on how efficacy of these compounds is altered by stress-induced pathology.

It is widely accepted that 5-HT_4_Rs stimulate cAMP production (65,66), indicating an interaction with the G_s_ protein pathway (67,69). Somewhat unexpectedly, in our signaling assays, PF-04995274 and other effective novel compounds behaved as inverse agonists or neutral antagonists – decreasing 5HT-induced activity below or to baseline levels. Within the HPC, cAMP signaling pathways are crucial for long term potentiation (LTP) and ultimately, memory consolidation (71,72). In a mouse model of chronic unpredictable stress (CUS), increased behavioral despair was accompanied by a downregulation of cAMP signaling (73) while increasing cAMP signaling improved behavioral responses to stress (74). Although we could not characterize *in vivo* activity of 5-HT_4_Rs, inverse agonism or neutral antagonism may lead to a reduction of receptor activity, perhaps by competition with endogenous 5-HT. This reduction in activity in neural circuits may counterintuitively underlie drug efficacy.

It is also possible that signaling pathways other than G_s_ and cAMP may play a critical role in the behavioral effects we have observed. Recent work has documented 5-HT_4_R interactions with other G-proteins, including G_i_ (75,76). 5-HT_4_Rs also recruit arrestin (77) and activate the extracellular signal-regulated kinase (ERK) pathway through interaction with the non-receptor tyrosine kinase Src, independent of G-protein signaling (78–80). In addition to influencing cellular differentiation and growth (81,82), activation of the ERK pathway is implicated in synaptic plasticity. Pharmacologically inhibiting ERK signaling in CA1 impairs LTP (83), while inhibition in the lateral AMG impairs LTP and long-term memory of Pavlovian fear conditioning (84,85). Future work characterizing how PF-04995274 and novel compounds alter Src or G_i_ related pathways will be informative.

Our group previously described an electrophysiological signature following prophylactic administration of several compounds characterized by a reduction of large AMPAR-driven synaptic bursts in CA3 (18,50,51), characteristic of potentially sharp wave ripples (SPW-Rs). SPW-Rs are notable for their role in memory and can contribute emotional salience to events (86–88); elevated 5-HT can quell SPW-Rs (89). Here, prophylactic DL5 and DL7, which acted at G_s_ as neutral antagonists, and DL8, which acted as an inverse agonist, reduced mean amplitude of EPSCs and mean frequency of large AMPAR bursts in CA3. Future testing could determine if these compounds may reduce SPW-Rs. Although considered a NMDA antagonist, recent work suggests (*R,S*)-ketamine and its metabolites also influence AMPAR-mediated signaling, which may contribute to antidepressant efficacy (18,90–93). It is possible that inverse agonists may impact AMPAR signaling by influencing cAMP-PKA or other ERK signaling pathways. Some work suggests that modulators of AMPAR signaling increase BDNF, which underlies antidepressant efficacy (94,95), but recent work failed to show a robust relationship between 5-HT_4_R activity and BDNF (96). Future work will elucidate if these compounds similarly alter activity in other brain regions.

Following a test of behavioral despair, PF-04995274, DL5, DL7, and DL8 increased neural activity in the DG, while PF-04995274, DL5 and DL7 increased activity in CA3 as measured by c-Fos^+^. We previously identified that prophylactic (*R,S*)-ketamine increases ΔFosB following social defeat (56) and c-Fos expression following CFC (51) in vCA3. Additionally, prophylactic (*R,S*)-ketamine also reduces large AMPAR-driven synaptic bursts in CA3 (18,50,51). Here, DL5 and DL7 exhibit a similar electrophysiological signature while also increasing c-Fos expression in vCA3. These seemingly contradictory results may be a common signature across compounds effective at improving stress-induced maladaptive behaviors. Within the HPC, 5-HT_4_Rs are expressed on excitatory (19) and GABAergic neurons (97,98), as well as astrocytes (104), making it likely that compounds targeting 5-HT_4_Rs in this region influence activity in a complex manner. It is possible that dampening activation of 5-HT_4_Rs via inverse agonism on interneurons ultimately disinhibits neural circuits underlying depressive-like behavior. Previous work has shown that 5-HT_4_R activation can modulate GABAergic activity within the HPC and other regions in a complex, bidirectional manner (94–96). Future work will explore how these compounds influence neurotransmission within the HPC.

The potential for 5-HT_4_R compounds to improve behavior in aged mice is promising. In older populations, treatment options for late-life depression (LLD) are lacking (102,103,105) as treatments such as SSRIs and esketamine show reduced efficacy (59,60,62). As LLD is a modifiable risk factor for cognitive impairment, (106–108) new therapeutics are needed. Unlike other 5-HT receptors that exhibit reduced binding with age, 5-HT_4_R expression is stable (109–111) making them a promising therapeutic target for LLD. Our group previously found that prophylactic (*R,S*)-ketamine is not effective in aged mice (57,58), but here we report that RS-67,333, PF-04995274, and DL7 are effective at reducing behavioral despair. 5-HT_4_R agonists may also improve cognitive function. Previous work found RS-67,333 improved object recognition in aged rats (110), and clinically, a subchronic, low-dose of prucalopride improved cognitive function in an image recall task (37) and enhanced connectivity between the central executive network and cingulate cortex (38). Future work will be needed to determine if acute or subchronic administration of the neutral antagonist DL7 may improve cognition.

We identified sex differences in how 5-HT_4_-targeting compounds influence behavior. In males, prophylactic administration of novel compounds improved behavioral despair, hyponeophagia, and perseverative behavior, but in females, efficacy is observed in reductions in hyponeophagia and perseverative behavior (**Table S1**). In males, compounds administered after stress reduce behavioral despair; in females compounds reduce hyponeophagia (**Table S2**). Although sex differences in 5-HT signaling have been documented (113–115) differences in 5-HT_4_Rs are underexplored. Post-mortem studies did not identify sex differences in 5-HT_4_R density (25,114); however, PET imaging identified lower 5-HT_4_R binding in the HPC (117) and limbic regions (109) of women compared to men. Hormone status may also influence 5-HT_4_R activity; in healthy men, testosterone level was negatively correlated with 5-HT_4_R binding, particularly in the HPC (118). In women, oral contraception use resulted in a 12% reduction in global 5-HT_4_R binding compared to controls (119). If hormone status decreases binding due to a decline in expression of the 5-HT_4_R or altered serotoninergic tone remains to be elucidated.

5-HT neurotransmission, and 5-HT_4_R activity, has been implicated in feeding behaviors. Increased 5-HT tone is anorexiogenic (120,121). Stimulating 5-HT_4_Rs in the nucleus accumbens induces hypophagia (122); conversely, inactivating 5-HT_4_Rs in this region increases feeding (122). 5-HT_4_R knock-out (KO) mice do not exhibit stress-induced hypophagia (123). Given their role in feeding behaviors, 5-HT_4_Rs have been implicated in anorexia nervosa restricting-type (AN) (124,125), a disease marked by restricted food intake (126–129). Although we did not test these compounds in a mouse model of AN, we find that following a brief period of food restriction, administration of inverse agonists PF-04995274 or DL8, as well as the neutral antagonist DL5, improved feeding behavior in the NSF in both sexes. We hypothesize that by blunting 5-HT_4_R activity, these compounds may protect against stress-induced hyponeophagia. Future testing using an activity-based anorexia model (130) may provide valuable insight into efficacy of compounds to improve AN.

In untreated patients with MDD, there is a 7% reduction in global 5-HT_4_R binding compared to healthy controls (131). There is also a negative correlation between global 5-HT_4_R binding and anxiety symptoms in patients with MDD, with more severe anxiety associated with lower 5-HT_4_R binding (132). In the U.S., only two 5-HT_4_R agonists are FDA-approved, tegaserod and prucalopride, for gastrointestinal disorders; (35,36,133) no 5-HT_4_R therapeutics are approved for psychiatric use. As we have demonstrated that a single administration of novel, small molecules targeting the 5-HT_4_R is effective, future clinical work might utilize acute or subchronic administration, reducing the likelihood of peripheral effects while maintaining efficacy. Additional positive findings from future metabolic and safety tests will allow these compounds to move towards clinical trials and may ultimately expand treatment options for those living with psychiatric illness.

## ACKNOWLEDGEMENTS AND DISCLOSURES

We thank Dr. Ronald Katz for initial assistance with funding and executing this design. We thank Dr. Nicole Perry-Hauser for construction of the CRE reporter construct. Figures of behavioral timelines were created with BioRender.com.

## AUTHOR CONTRIBUTIONS

S-XD, and DWL designed the compounds. RR, BKC, CAD, IMD, DJD, and JAJ designed research experiments. RR and CAD wrote the manuscript, which was edited by all the authors. RR, BKC, and MA performed behavioral research and analysis. MA performed immunohistochemistry and cell counting. MN performed the cell-based assays shown, with preliminary experiments by MP. VL and GT conducted electrophysiology experiments.

## ADDITIONAL INFORMATION

### Supplementary Information

The online version contains supplementary material available at X.

### Address correspondence

to Briana K. Chen, Ph.D., at or Christine A. Denny, Ph.D., at.

### Reprints and permission information

is available at X.

## SUPPLEMENTAL INFORMATION

### SUPPLEMENTAL METHODS

#### Drugs

All drug injections were administered intraperitoneally (i.p.) in volumes of 0.1 cc per 10 mg body weight. PF-04995274 (Sigma, Catalog No 1331782-27-4, St. Lois, MO), prucalopride (Sigma, Catalog No 179474-8108, St. Lois MO), and RS-67,333 (Sigma, Catalog No 168986-60-5, St.

Lois, MO) were dissolved in a 5% dimethyl sulfoxide (DMSO)-0.9% NaCl (saline) solution (BioLogics, Model 3000, Manassas, VA). Novel 5-HT_4_R compounds would not remain in a 5% DMSO solution; instead, compounds were dissolved in a 10% 2-hydroxypropyl-beta-cyclodextrin (HBCD) solution (1,2). All drugs were administered in a single injection either 1 week before stress or 5 min after stress at 3 or 10 mg/kg of body weight based on our previous work (3) All drugs and materials are listed in **Supplemental Table S3**.

#### Molecular Assays

##### Materials

HEK-293T cells were obtained from the American Type Culture Collection (Rockville, MD) and were cultured in a 5% CO_2_ atmosphere at 37°C in Dulbecco’s Modified Eagle Medium (DMEM) (Life Technologies, Grand Island, NY) supplemented with 10% Fetal Bovine Serum (FBS) and 100 IU ml^−1^ penicillin and 100 μg ml^−1^ streptomycin (Corning, Tewksbury, MA). The following chemicals were used without further modification: Serotonin (Tocris, Bristol, United Kingdom), Coelenterazine H (Dalton Pharma Services, Toronto, ON, Canada), Polyethylenimine (PEI; Polysciences, Warrington, PA), Firefly D-luciferin (Nanolight Technologies, Norman, OK), Prucalopride (Sigma, Catalog No. 179474-8108, St. Louis, MO), RS-67,333 (Sigma, Catalog No. 168986-60-5, St. Louis, MO), and PF-04995274 (Sigma, Catalog No. 1331782-27-4, St. Louis, MO).

##### DNA Constructs

The vector coding for 5-HT_4b_ was purchased from cDNA.org and utilized in miniG recruitment experiments. 5-HT_4c_ was created by modifying the 5-HT_4b_ cDNA using standard procedures and was sequence verified (Plasmidsaurus, Eugene, OR) and 5-HT_4c_ was utilized in cAMP experiments. The plasmids coding for CAMYEL, Venus-miniGs and 5-HT_4b_-nanoluc (5-HT_4b_-nluc) were gifts from Dr. Nevin Lambert at the Medical College of Georgia. Full sequences of both isoforms are listed in **Supplemental Table 4**.

##### Transfection

For the miniGs or the CAMYEL BRET assays, a total of 1, 5, or 20 µg of cDNA, respectively, was transiently transfected into HEK-293T cells (350,000 cells for CRE in 12-well plate format; 2 × 10^6^ cells per plate for miniGs and 4 × 10^6^ cells per plate for CAMYEL and arrestin in 10 cm dishes). For the miniGs assay, 1 µg receptor-nluc, 1 µg Venus-miniGs, and 3 µg empty vector were transfected using PEI in a 6:1 ratio (diluted in DMEM). For the CAMYEL assay, 1.25 µg of receptor, 10 µg of CAMYEL sensor, and 8.75 µg of empty vector were transiently transfected using PEI in a 2:1 ratio (diluted in DMEM). Cells were maintained and transfected in the HEK-293T media described above, then switched to a serum free version of the media (lacking Fetal Bovine Serum (FBS)). For the CAMYEL and miniGs assay, media was switched to serum free 2-6 hours post transfection. CAMYEL and miniGs assays were performed 24 hours after transfection.

##### Bioluminescence Resonance Energy Transfer (BRET)

Experiments were performed as described previously (4). Briefly, transfected cells were dissociated and resuspended in Dulbecco’s Phosphate Buffered Saline (dPBS). Cells were added to a black-framed, white well 96-well plate (Perkin Elmer, Waltham, MA). The luciferase substrate coelenterazine H (5 µM) was added to each well at time zero. Five min later, ligands were added. After a 10 min incubation, the BRET signal was measured. BRET measurements were performed using a PHERAstar FS plate reader or a LUMIstar plate reader (BMG Labtech, Cary, NC). The BRET signal was calculated as the ratio of the light emitted by the acceptor (510–540 nm) over the light emitted by the donor (475 nm). Dose–response curves were fit using a three-parameter logistic equation in GraphPad Prism 10 (La Jolla, CA). All experiments were repeated in at least three independent trials each with triplicate determinations.

#### Behavioral Assays

##### Contextual Fear Conditioning (CFC)

A 3-shock CFC was administered as previously described (3,5). CFC was conducted in chambers obtained from Actimetrics (Lafayette, IN), with internal dimensions of 7.4^“^ L x 8.1^”^ D x 7.9^”^ H. The chambers had metal walls on each side, clear plastic front and back walls and ceilings, and stainless-steel bars on the floor. A house light (CM1820 bulb, 28v, 100mA) mounted directly above the chamber provided illumination. Each chamber was located inside a larger, insulated, plastic cabinet that provided protection from outside light and noise. Each cabinet contained a ventilation fan that was operated during the sessions. A paper towel dabbed with lemon solution (Pure Lemon Extract, McCormick®, Hunt Valley, MD) was placed underneath the chamber floor. To regulate the amount of lemon solution, Q-Tips were used to place 4 equal dots of lemon scent on 4 corners of the paper towel lining the chamber. Mice were held outside the experimental room in their home cages prior to testing and transported to the conditioning apparatus individually in standard mouse cages. Chambers were cleaned with 70% EtOH after each run. Mice were placed into the conditioning chamber and received shocks at 180 s, 240 s, and 300 s (2 s duration, 0.75 mA). Fifteen seconds after the last shock, mice were removed from the chamber. Overall, the training session lasted 317 s. During re-exposure, mice were placed in the conditioning chamber for 5 minutes and did not receive any shocks. All sessions were scored for freezing using FreezeFrame4.

##### Learned Helplessness (LH)

LH was used to induce depressive-like behavior through exposure to repeated unpredictable foot shocks and as previously described (6). In this paradigm, mice are exposed to unpredictable and uncontrollable stress (shocks) and then develop coping deficits to deal with the inescapable shocks. We utilized a two-chamber shuttle box (ENV 010MD; Med Associates, St. Albans VT) within a sound-attenuated cubicle. The grid floor was made of stainless steel and connected to a shock generator. The scrambled shock generator (ENV 414S, Med Associates, St. Albans VT) created varying electrical potential differences between bars preventing an animal from avoiding shock.

*Inescapable shock (training):* For each shuttle box, 2 animals were administered the protocol at the same time; the central door was closed, with one animal in the chamber on each side. After a 3 min habituation period, the shock deliveries began. The training protocol consisted of 70 shocks, each with a 3 s average duration, at 0.5 mA, and with an intertrial interval (ITI) of approximately 15 s.

*Shock escape (testing):* Mice were tested in the same shuttle box used in the inescapable shock training. The box consisted of two identical chambers (17 l x 20 w x 17 h), separated by an automated door that opened vertically. The shuttle box was equipped with 8 infrared beams (4 on each side) for detecting position and activity of the animal (Med Associates, St. Albans, VT). Five days after training, each mouse was placed into the right chamber with the door raised and was allowed to freely explore both chambers for 3 min. Then the door then closed automatically. At the beginning of each trial, the door was raised and 5 s later a foot shock (0.5 mA) was delivered. The subject’s exit from the shocked side ended the trial. If the mouse did not exit after 15 s, the shock was turned off and the trial ended. The door was lowered at the end of the trial. A session consisted of 30 trials, each separated by a 30 s ITI. Escape latencies were computed as the time from shock onset to the end of trial. If the subject failed to make a transition the maximum 15 s was used for the escape latency score.

##### Forced Swim Test (FST)

The FST was administered as previously described (3,6) to gauge behavioral despair. Briefly, mice were placed into clear plastic buckets 20 cm in diameter and 23 cm deep filled 2/3^rd^ of the way with 22°C water. Mice were recorded from the side for 6 min and were exposed to the test on 2 consecutive days. Immobility was scored by an experimenter blind to experiment group and was used as a measure of depressive behavior.

##### Marble Burying (MB)

The MB assay was conducted in a clean cage (10.5 in x 5.5 in) containing soft pliable Beta Chip bedding (Northeastern Products Corp, Warrensburg, NY). The cage contained 16 marbles set up in 4 rows of 4 across. Mice were given 30 minutes to explore and bury. Following task completion, the percentage of marbles buried was calculated, a marble was counted as buried if at least if 2/3^rd^ was covered.

##### Novelty-Suppressed Feeding (NSF)

Testing was performed as previously described (6–8). Briefly, the NSF testing apparatus consisted of a plastic box (50 x 50 x 20 cm). The floor of which was covered with approximately 2 cm of wooden bedding and the arena was brightly lit. Mice were food restricted for 12 h prior to testing. At the time of testing, a single pellet of food (regular chow) was placed on a white paper platform positioned in the center of the box. Each animal was placed in a corner of the box, and a stopwatch was immediately started. The latency of the mice to begin eating in the arena was recorded. Immediately after the latency was recorded, the food pellet was removed from the arena. The mice were then placed back into their home cage. The latency to eat and the amount of food consumed in 5 min were measured (home cage consumption), followed by an assessment of post-restriction weight. A Kaplan-Meier survival analysis was used due to the lack of normal distribution of data. The Mantel-Cox log-rank test was used to evaluate differences between the experimental groups.

##### Open Field (OF)

The OF assay utilized eight plexiglass open field boxes (35 x 35 x 38 cm) equipped with a SmartFrame 16X-16Y infrared beam system (Kinder Scientific, Chula Vista, CA). All x-y ambulatory movements and rearing and zone metrics were captured from beam breaks using MotorMonitor (v 5.05, Kinder Scientific). The system defined the center and surround regions, with the center zone covering a quarter of the total square area (17.4 x 17.4 cm) in the center of the apparatus approximately 8.7 cm from the perimeter.

#### Electrophysiology

Slice electrophysiology was conducted as previously described (3, 7–9). One week after saline, PF-0499527, or novel 5-HT_4_R agonists administration (3 mg/kg) mice were anesthetized by isoflurane inhalation, decapitated, and brains rapidly removed. CA3 slices (350 µm) were cut on a vibratome (Leica VT1000S) in ice cold partial sucrose artificial cerebrospinal fluid (aCSF) solution (in mM): 80 NaCl, 3.5 KCl, 4.5 MgSO_4_, 0.5 CaCl_2_, 1.25 H_2_PO_4_, 25 NaHCO_3_, 10 glucose, and 90 sucrose equilibrated with 95% O_2_/5% CO_2_ and stored in the same solution at 37°C for 30 min, then at RT until use. Recordings were made at 30 – 32°C (TC234-B, Warner Instrument Corp.) in aCSF (in mM: 124 NaCl, 8.5 KCl, 1 NaH_2_PO_4_, 25 NaHCO_3_, 20 glucose, 1 MgCl_2_, 2 CaCl_2_). Whole-cell voltage clamp recordings (-70 mV) were obtained using a patch pipette (4-6 M MΩ) containing in mM: 135 K-Gluconate, 5 KCl, 0.1 EGTA-Na, 10 HEPES, 2 NaCl, 5 ATP, 0.4

GTP, 10 phosphocreatine (pH 7.2; 280 – 290 mOsm). Bicuculline (5 µM) was also included in the bath solution to inhibit GABA_A_ receptors. Patch pipettes were made from borosilicate glass (A-M Systems, Sequium, WA) using a micropipette puller (Model P-1000, Sutter Instruments). Recordings were made without correction for junction potentials. Pyramidal cells were visualized and targeted via infrared-differential interference contrast (IR-DIC, 40x objective) optics on an Axioskop-2 FS (Zeiss, Oberkochen, Germany). Data was exported from AxoGraph (v 1.4.4) into Excel for analysis. Briefly, the baseline period was subtracted and the signal filtered (100 Hz). Amplitude values of all negative peak data (> 10 pA) was exported with a decay of 90% - 10%. Data was sorted by absolute value and burst activity defined as an absolute value greater than 100 pA.

#### Immunohistochemistry

Immunohistochemistry was performed as previously described (2,5). Mice were deeply anesthetized with a ketamine/xylazine mixture 1 hour following FST and were transcardially perfused with 50 mL ice-cold 1x PBS followed by 50 mL 4% paraformaldehyde (PFA). Brains were removed and left in 4% PFA overnight and then cut on a vibratome (Leica VT1000S, Deer Park, IL, USA) and stored in 1x PBS with 0.1% sodium azide prior to processing. Sections (50 µm) were washed 3x in 1x PBS and blocked in 1x PBS with 0.5% Triton X-100 (PBST) and 10% normal donkey serum (NDS) for 2 hours at room temperature (RT). Sections were then incubated in primary rat anti-c-Fos for 24 hr at 4°C (Synaptic Systems, 226 017, 1:5,000). The next day tissue was washed 3x in 1x PBS and then incubated for 2 hours in secondary donkey anti-rat Alexa Fluor 488 at RT (ThermoFisher, A21208, 1:500). Tissue was incubated with Hoechst in 1x PBS at RT (Hoechst, 33342, ThermoFisher, 1:10k) and washed again in 1x PBS. All slices were mounted on glass slides and cover slipped with Fluoromount G (Electron Microscopy Sciences, Hatfield, PA, 17984-25).

#### Confocal Microscopy

Fluorescent confocal micrographs were taken at 20x magnification with a Leica SP8 microscope and with LAS X software as previously described (7). Hippocampal sections were imaged throughout the rostro-caudal axis of the HPC. Identification of hippocampal regions (CA1, CA3, and DG) involved acquiring 5 – 8 sections per mouse due to variability in sectioning. Images were scanned and captured at a z-increment of 3 μm. Z-stack analysis was performed using LAS X image browser to determine expression of c-Fos which was compared across all sections using identical exposure conditions.

#### Cell Quantification

A researcher blinded to treatment group used FIJI software (NIH) to manually count c-Fos immunoreactive cells in the DG, CA3 and CA1 regions of the HPC of one hemisphere across the rostrocadual axis (-1.34 to -2.18 AP). The number of c-Fos^+^ cells across groups is presented in **Figure 6**.

#### Statistical Analysis

Results from all data analyses are expressed as means ± SEM. Alpha was set to 0.05. Data were analyzed using Prism v8.0 or 10.0 (GraphPad Software, La Jolla, CA). All experimental groups were analyzed together using one or two-way ANOVAs with repeated measures were applied to data as appropriate. Significant main effects and/or interactions were followed by Fisher’s LSD. All statistical tests and *p* values are included in **Supplemental Table S5.**

## SUPPLEMENTAL FIGURES AND FIGURE LEGENDS

**Figure S1.**
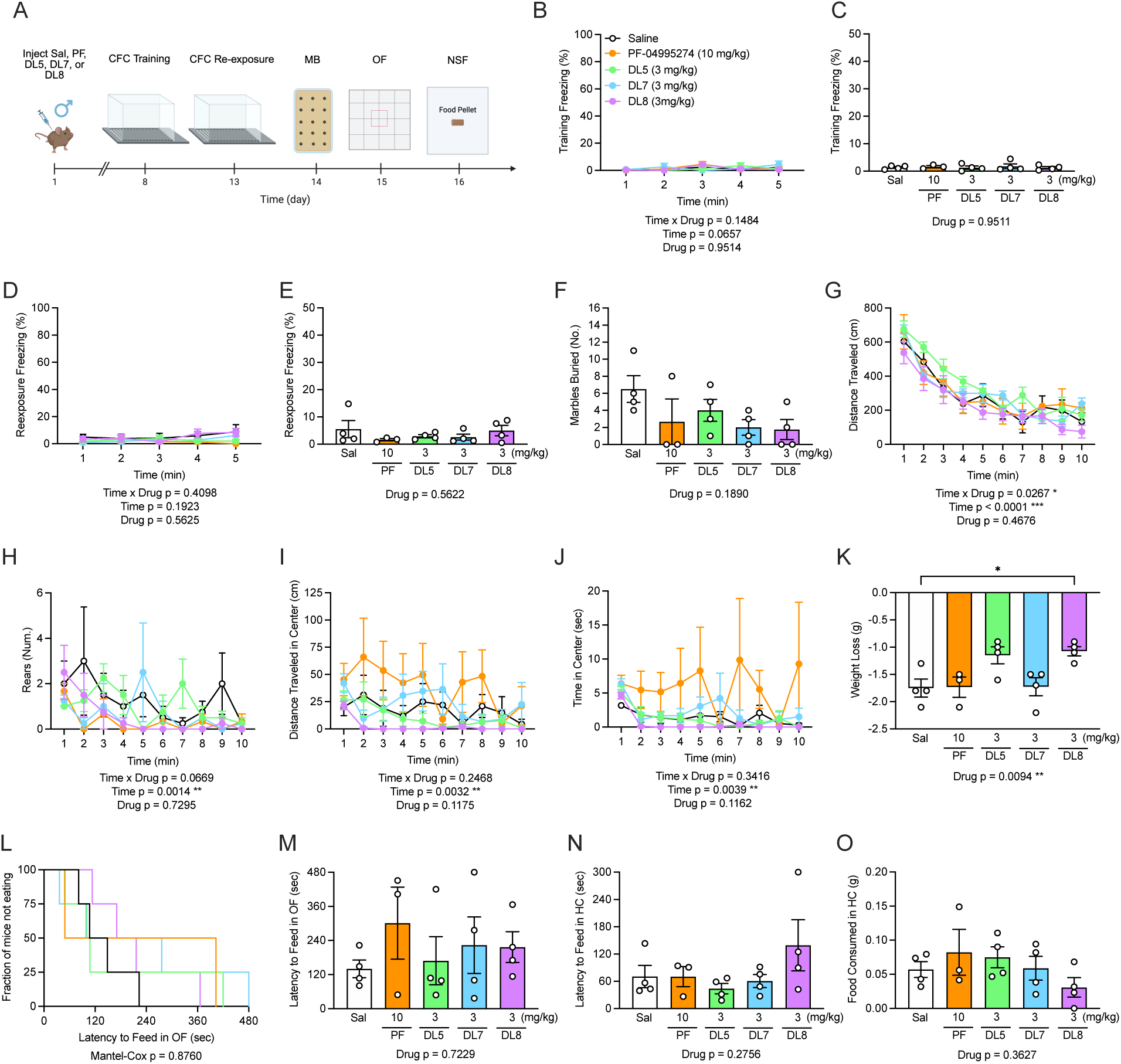
In the absence of stress, prophylactic administration of PF-04995274 or a novel compound does not influence behavior in male mice. **(A)** Experimental timeline. **(B, C)** Freezing across CFC no-shock training is comparable across groups. **(D, E)** During CFC exposure freezing is comparable across groups. **(F)** In the MB task, the number of marbles buried was similar across all drug groups. **(G, H)** In the OF, drug administration did not significantly alter the total distance traveled or number of rears compared to saline controls. **(I, J)** Distance traveled and time spent in the center zone of the OF was similar across all groups. **(K)** During the food restriction phase of NSF, mice administered DL8 (3 mg/kg) lost more weight compared to saline. **(L, M)** Latency to feed in the open arena in the NSF was comparable across groups. **(N)** Latency to feed in the home cage and **(O)** amount of food consumed was comparable across groups. Sal n = 4, PF n = 3, DL5 n = 4, DL7 n = 4, DL8 n = 4 male mice per group; error bars represent ± SEM; * p < 0.05, ** p < 0.01, **** p < 0.0001. CFC, contextual fear conditioning; MB, marble burying; NSF, novelty suppressed feeding; OF, open field; HC, home cage; Sal, saline; PF, PF-04995274; min, minutes; sec, seconds; mg, milligram; kg, kilogram. Behavioral timeline created with BioRender.com.

**Figure S2.**
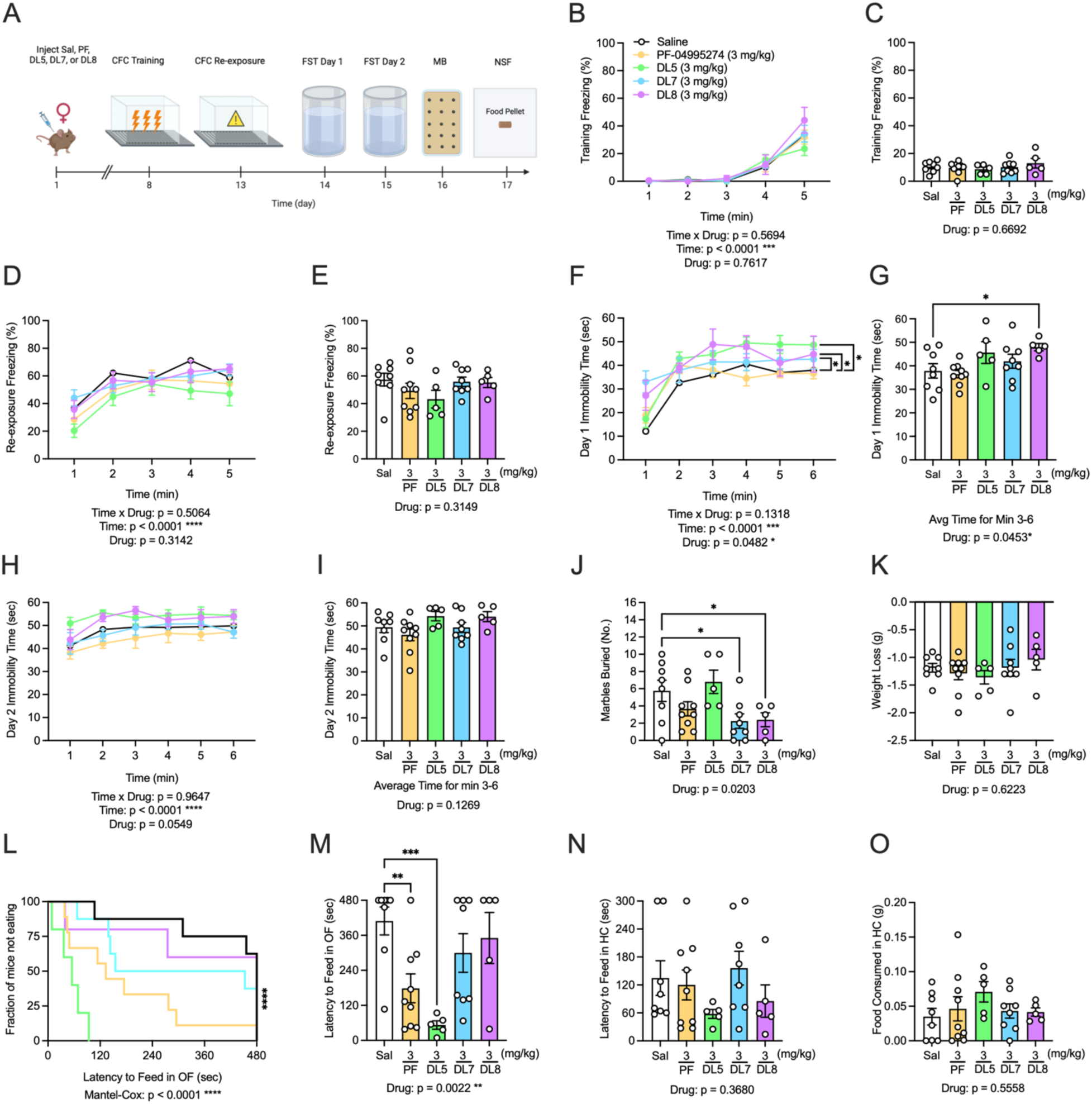
Prophylactic DL5 attenuates hyponeophagia while DL7 and DL8 attenuate perseverative behavior in adult female 129S6/SvEv mice. **(A)** Experimental protocol. **(B)** Freezing was comparable between all groups during CFC training. **(C)** Average freezing time was comparable between all groups during CFC training. **(D)** During context re-exposure, freezing was comparable between all groups. **(E)** Average freezing time was comparable between all groups during context re-exposure. **(F)** During day 1 of the FST, compounds DL5, D7, and DL8 (3 mg/kg) increased immobility compared to saline control across the entire test session, **(G)** but only DL8 (3 mg/kg) increased immobility compared to saline during minutes 3 – 6. **(H, I)** On day 2 of the FST, all groups exhibited comparable immobility time when compared with saline-administered mice. **(J)** In the MB task, D7 and DL8 (3 mg/kg) buried fewer marbles compared to saline control. **(K)** Weight loss was similar across groups during food restriction phase of NSF. **(L, M)** In the NSF, PF-04995274 and DL5 (3 mg/kg) exhibited reduced latency to feed in the OF. In the home cage, all groups exhibited comparable **(N)** latency to feed and **(O)** amount of food consumed. Sal n = 5, PF n = 9, DL5 n = 5, DL7 n = 8, DL8 = 5 female mice per group; error bars represent ± SEM; *p < 0.05, ** p < 0.01, *** p < 0.001. CFC, contextual fear conditioning; FST, forced swim test; HC, home cage; MB, marble burying; NSF, novelty-suppressed feeding; OF, open field; PF, PF-04995274; Sal, saline. Behavioral timeline created with BioRender.com.

**Figure S3.**
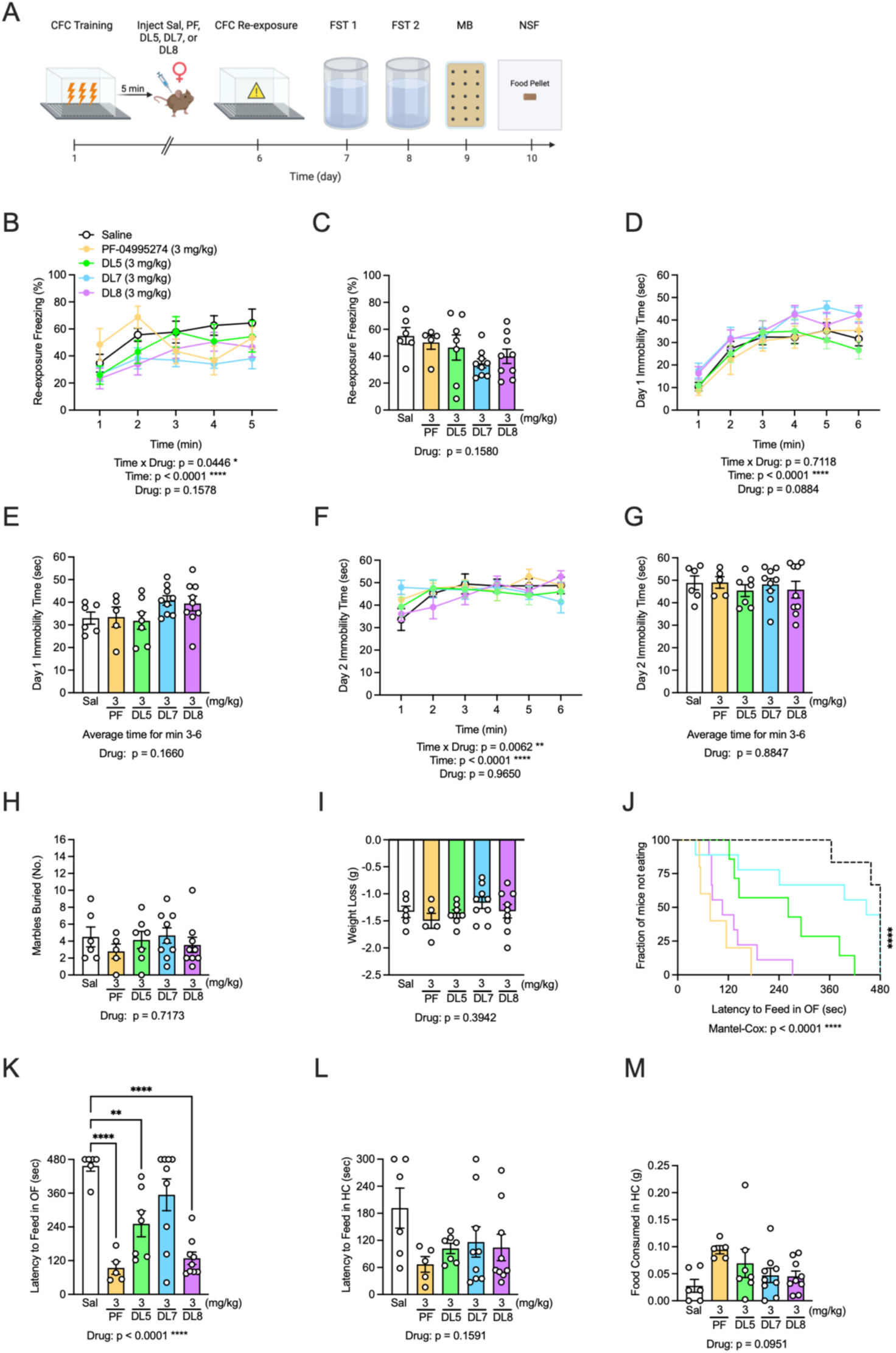
Post-stress administration of DL7 and DL8 decrease fear expression, while DL5 and DL8 decrease hyponeophagia in adult female 129S6/SvEv mice. **(A)** Experimental design. **(B, C)** Freezing was comparable across all groups during CFC exposure. **(D, E)** On day 1 of FST, all groups exhibited comparable immobility time. **(F, G)** On day 2 of FST all groups exhibited comparable immobility time. **(H)** In the MB task, all groups buried a comparable number of marbles. **(I)** In the NSF, all groups lost a comparable amount of weight during food restriction period. **(J, K)** PF-04995274, DL5, and DL8 (3 mg/kg) decreased the latency to feed in the open arena. **(L)** Latency to feed in the home cage was comparable across groups as was **(M)** the amount of food consumed in the home cage. Sal n = 6, PF n = 5, DL5 n = 7, DL7 n = 9, DL8 n = 9 female mice per group; error bars represent ± SEM; * p < 0.05, ** p < 0.01, *** p < 0.001. CFC, contextual fear conditioning; FST, forced swim test; MB, marble burying; NSF, novelty suppressed feeding; OF, open field; HC, home cage; Sal, saline; PF, PF-04995274; min, minutes; sec, seconds; mg, milligram; kg, kilogram. Behavioral timeline created with BioRender.com.

**Figure S4.**
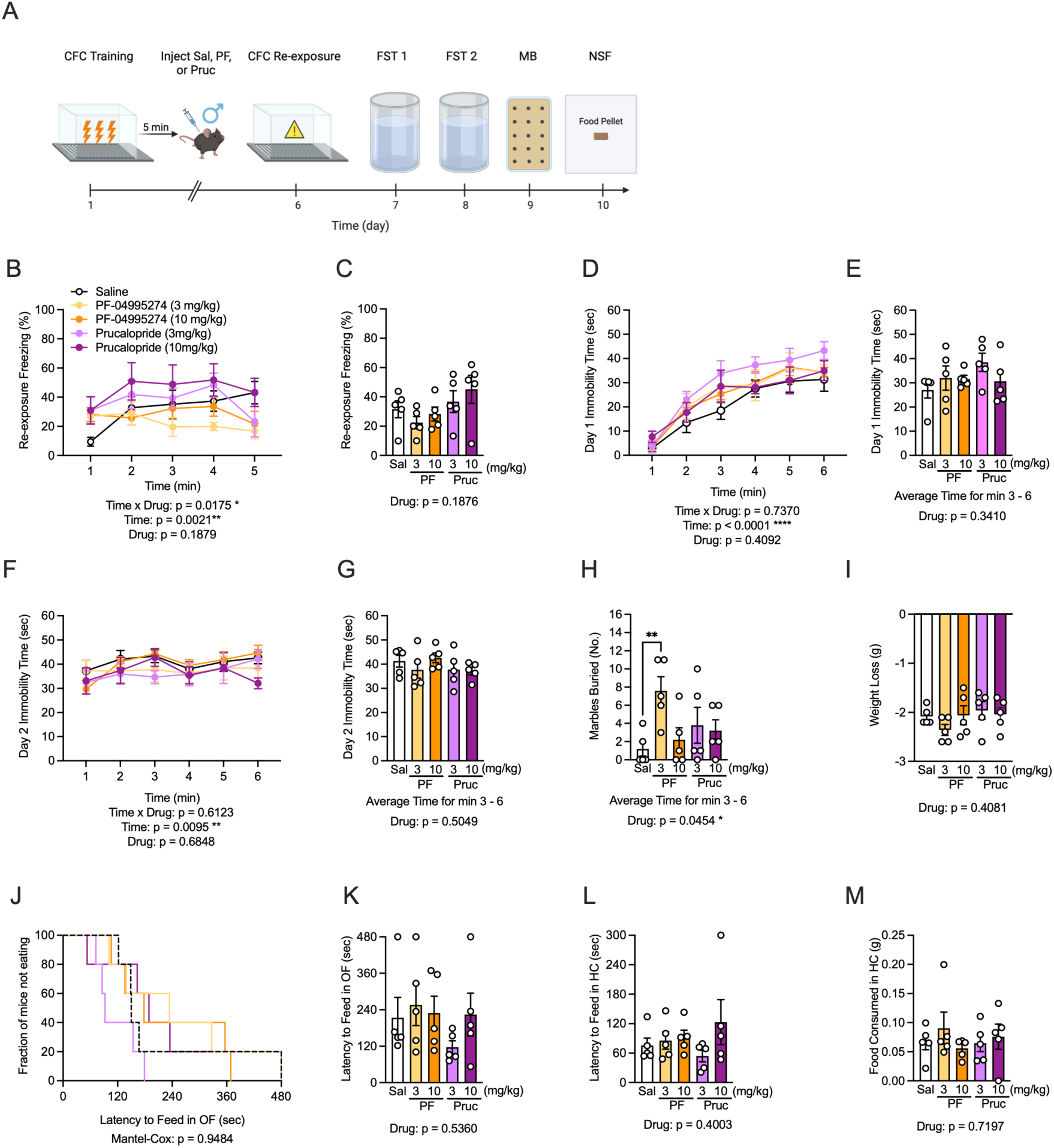
5-HT_4_R agonists do not improve stress induced maladaptive behavior when administered after stress in adult male C57BL/6J mice. **(A)** Experimental timeline. **(B, C)** Freezing across groups is comparable during CFC exposure. **(D, E)** On day 1 of FST and **(F, G)** day 2 of FST, immobility time was comparable across all groups. **(H)** In the MB task, PF-04995274 (3 mg/kg) increased the number of marbles buried compared to saline. **(I)** Weight loss during food restriction in the NSF is comparable across groups. **(J, K)** During the NSF latency to eat in the open arena was comparable across groups. **(L)** Latency to feed in the home cage and **(M)** food consumed in the home cage was comparable across groups. n = 5 male mice per all drug groups; error bars represent ± SEM; ** p < 0.01. CFC, contextual fear conditioning; FST, forced swim test; MB, marble burying; NSF, novelty suppressed feeding; OF, open field; HC, home cage; Sal, saline; PF, PF-04995274; Pruc, prucalopride; min, minutes; sec, seconds; mg, milligram; kg, kilogram. Behavioral timeline created with BioRender.com.

**Figure S5.**
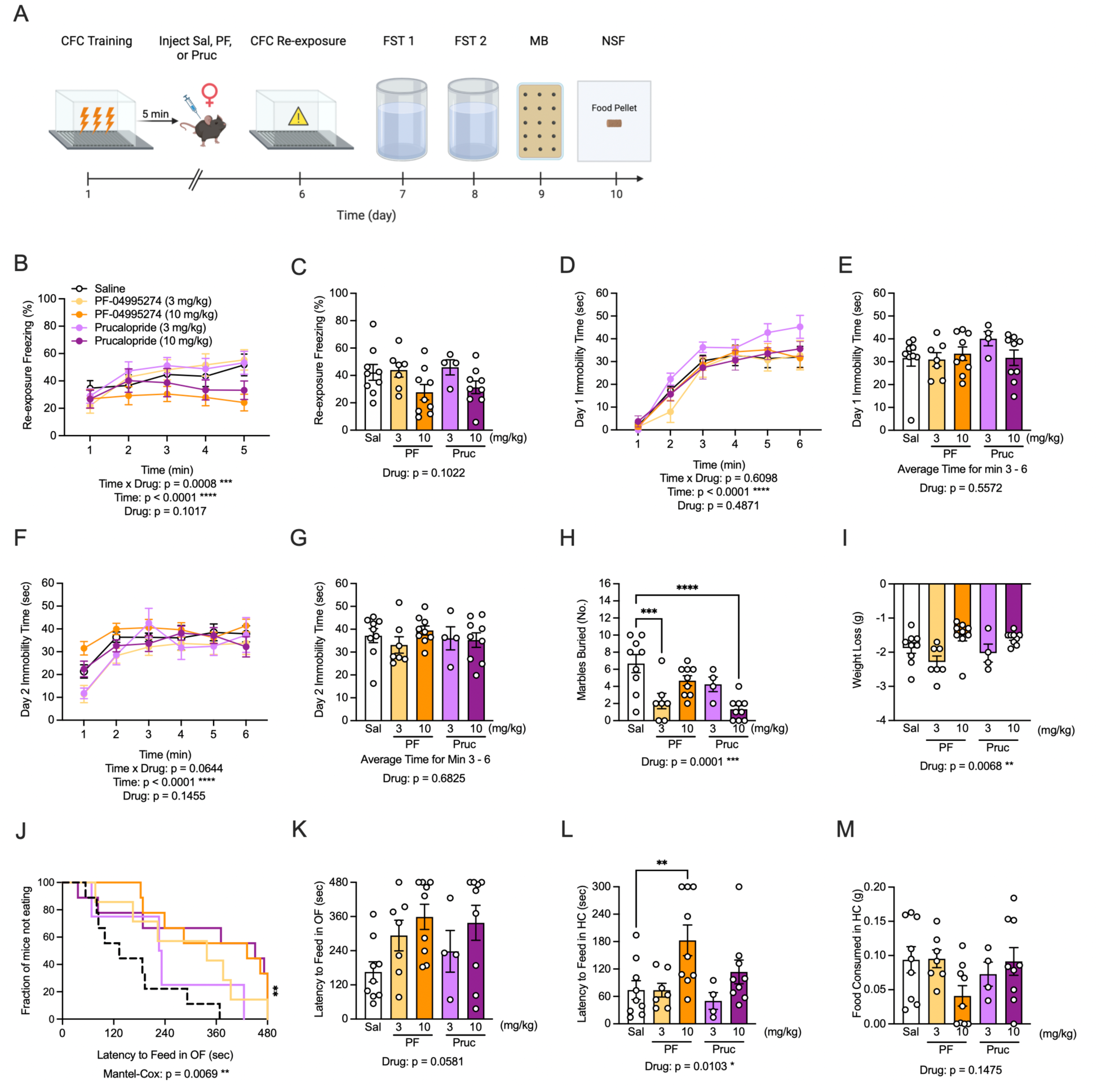
PF-04995274 and prucalopride improve perseverative behavior when administered after stress in adult female C57BL/6J mice. **(A)** Experimental timeline. **(B-C)** Freezing across groups is comparable during CFC exposure. **(D-E)** On day 1 of FST and **(F, G)** day 2 of FST immobility time was comparable across all groups. **(H)** In the MB task, PF-04995274 (3 mg/kg) and prucalopride (10 mg/kg) reduces the number of marbles buried compared to saline. **(I)** Weight loss during food restriction in the NSF across drug groups is comparable to saline controls. **(J, K)** During the NSF latency to eat in the open arena across drug groups was comparable to saline. **(L)** PF-04995274 (10 mg/kg) increased latency to feed in the home cage but **(M)** food consumed in the home cage was comparable across groups. Sal n = 9, 3 mg/kg PF n = 7, 10 mg/kg PF n = 9, 3 mg/kg Pruc n = 4, 10 mg/kg Pruc n = 9 female mice per group; error bars represent ± SEM; * p < 0.05, ** p < 0.01, *** p < 0.001. CFC, contextual fear conditioning; FST, forced swim test; MB, marble burying; NSF, novelty suppressed feeding; OF, open field; HC, home cage; Sal, saline; PF, PF-04995274; Pruc, prucalopride; min, minutes; sec, seconds; mg, milligram; kg, kilogram. Behavioral timeline created with BioRender.com.

**Figure S6.**
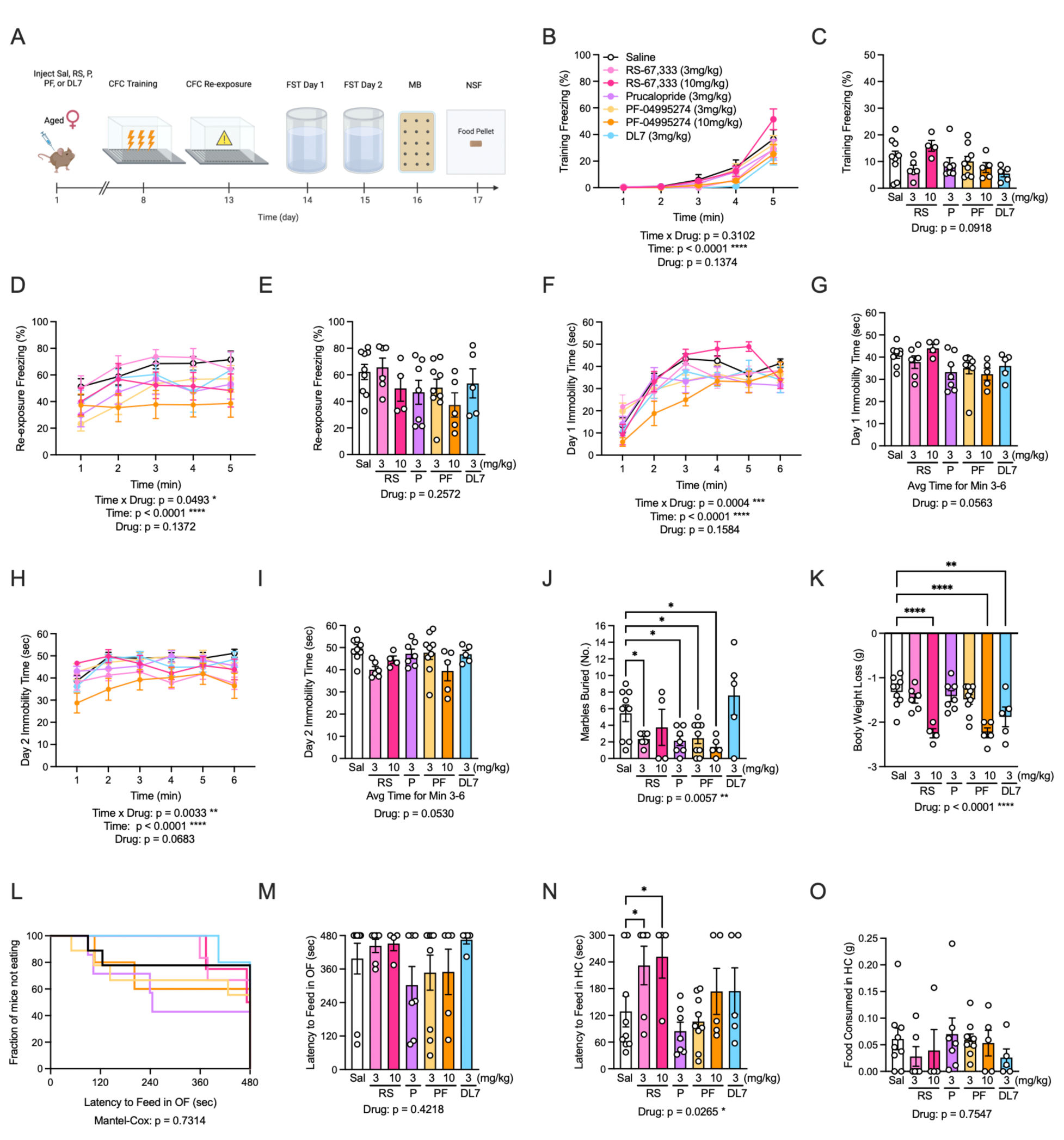
Prophylactic RS-67,333, prucalopride, and PF-04995274 attenuate perseverative behavior in aged female 129S6/SvEv mice. **(A)** Experimental timeline for aged (∼17 month) female mice. **(B, C)** Freezing across CFC training is comparable across groups. **(D, E)** During CFC exposure freezing is comparable across groups. **(F, G)** On day 1 and **(H, I)** day 2 of FST immobility was comparable across groups. **(J)** In the MB task, RS-67,333 (3 mg/kg), prucalopride (3 mg/kg), and PF-04995274 (3 and 10 mg/kg) reduced the number of marbles buried compared to saline. **(K)** During the food restriction phase of NSF, RS-67,333 (10 mg/kg), PF-04995274 (10 mg/kg) and DL7 (3 mg/kg) groups lost more weight compared to saline. **(L, M)** Latency to feed in the open arena in the NSF was comparable across groups. **(N)** RS-67,333 (3 and 10 mg/kg) increased latency to feed in the home cage compared to saline but **(O)** amount of food consumed was comparable across all groups. Sal n = 9, 10 mg/kg RS n = 6, 3 mg/kg RS = 4, Pruc n = 7, 3 mg/kg PF n = 9, 10 mg/kg PF = 5, DL7 n = 5 female mice per group; error bars represent ± SEM; * p < 0.05, ** p < 0.01, **** p < 0.0001. CFC, contextual fear conditioning; FST, forced swim test; MB, marble burying; NSF, novelty suppressed feeding; OF, open field; HC, home cage; Sal, saline; PF, PF-04995274; P, prucalopride; RS, RS-67,333; min, minutes; sec, seconds; mg, milligram; kg, kilogram. Behavioral timeline created with BioRender.com.

## SUPPLEMENTAL TABLES

**Table S1.**
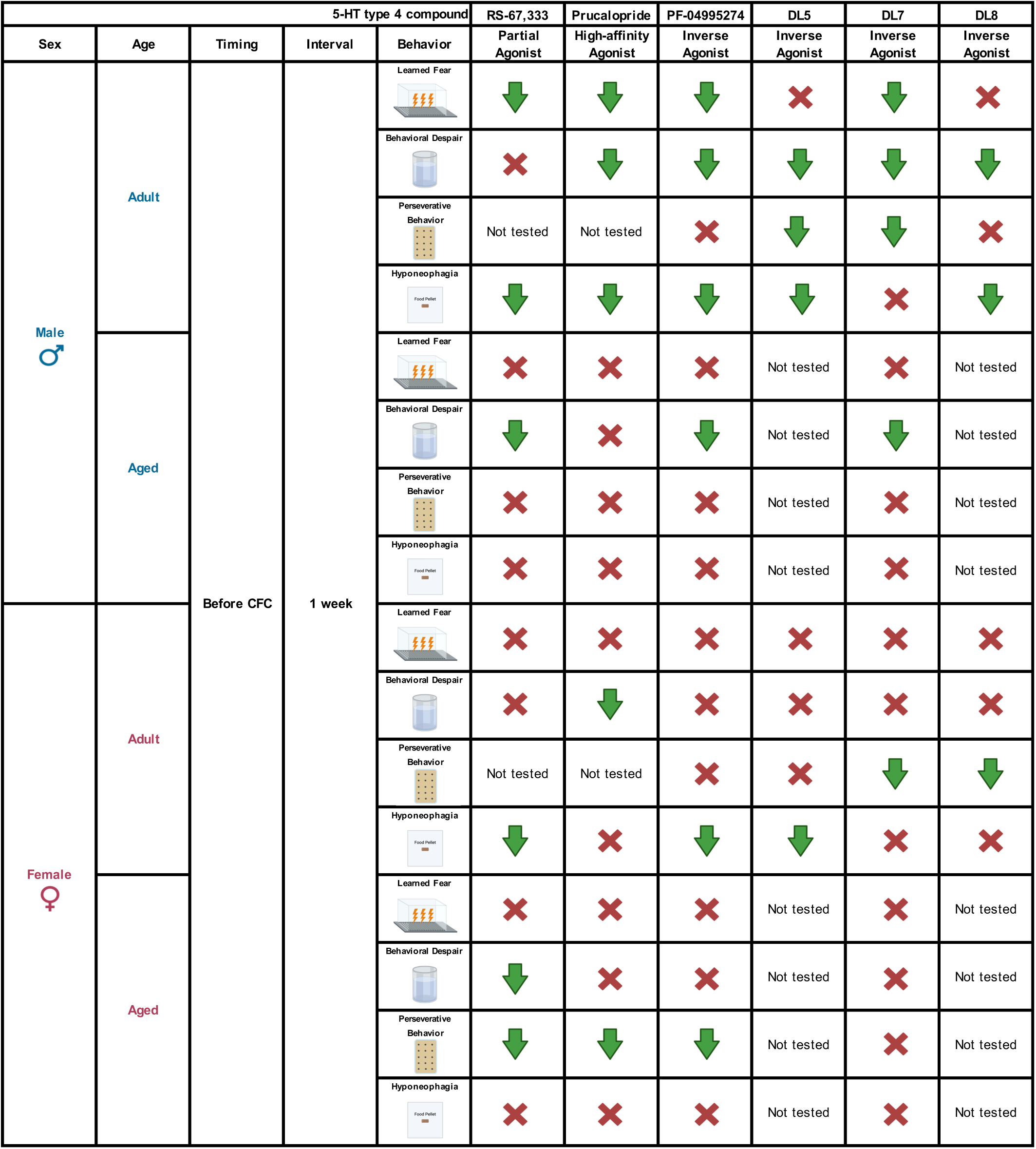
Prophylactic behavioral summary.

**Table S2.**
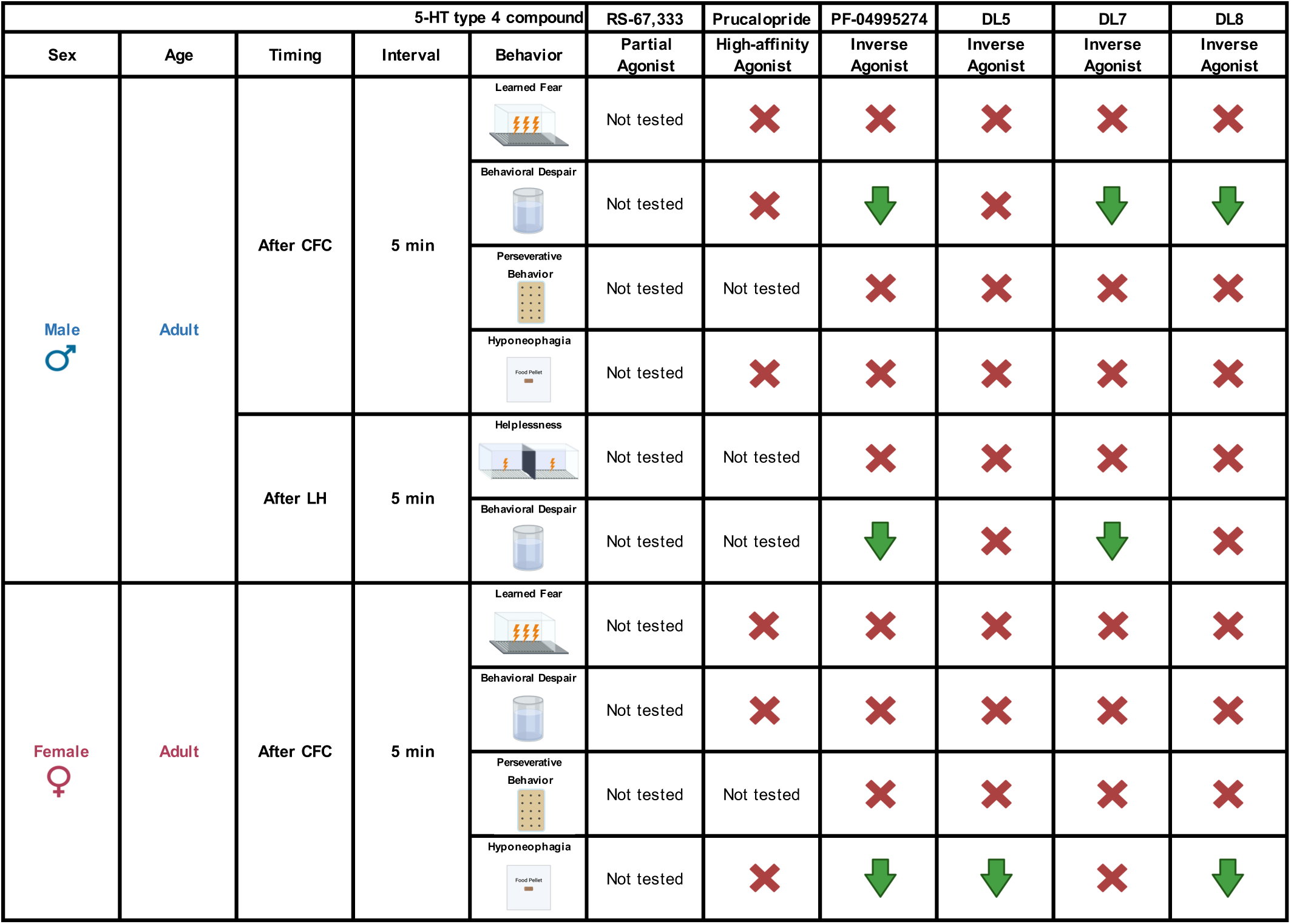
Post-stress behavioral summary.

**Table S3.**
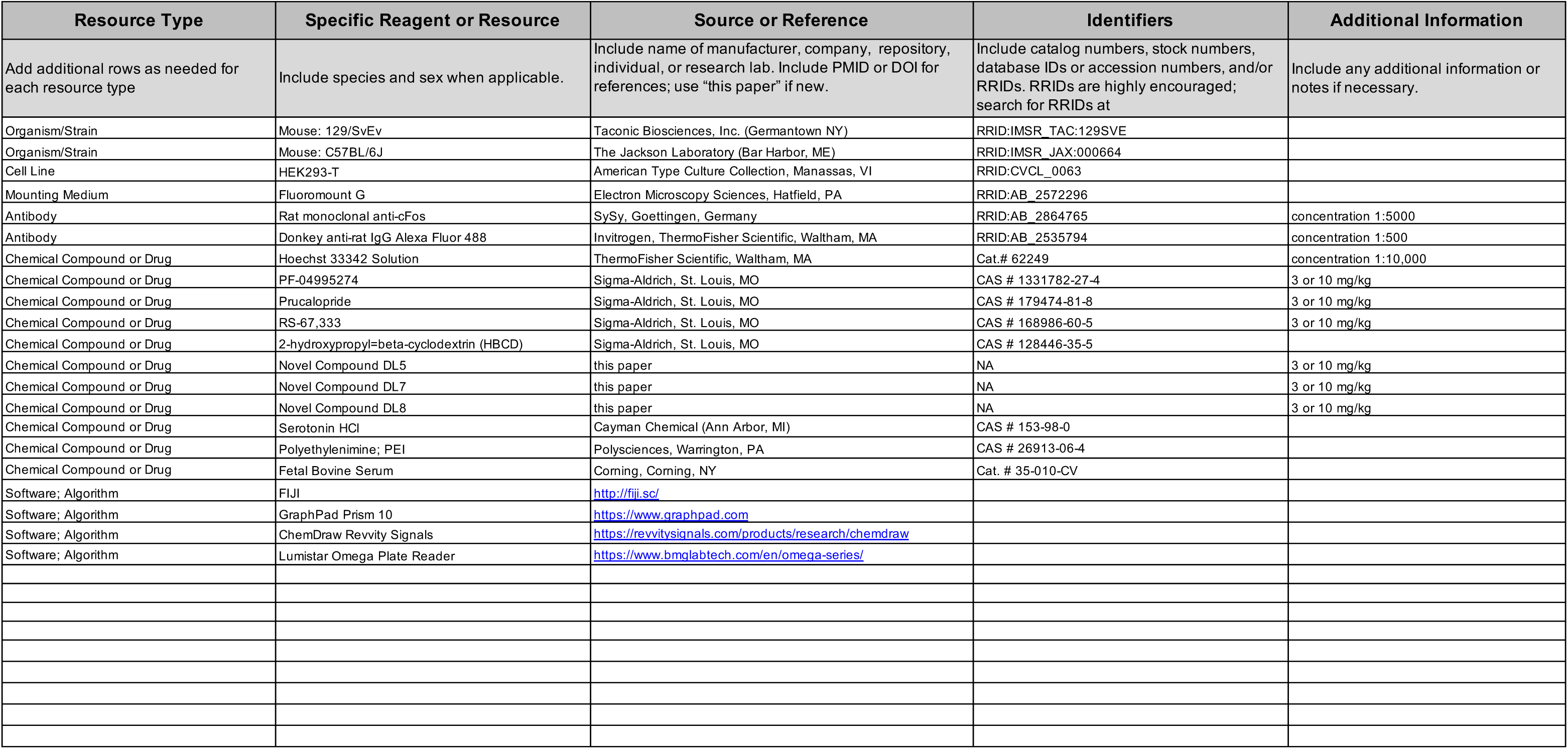
Key Resources Table.

**Table S4.**
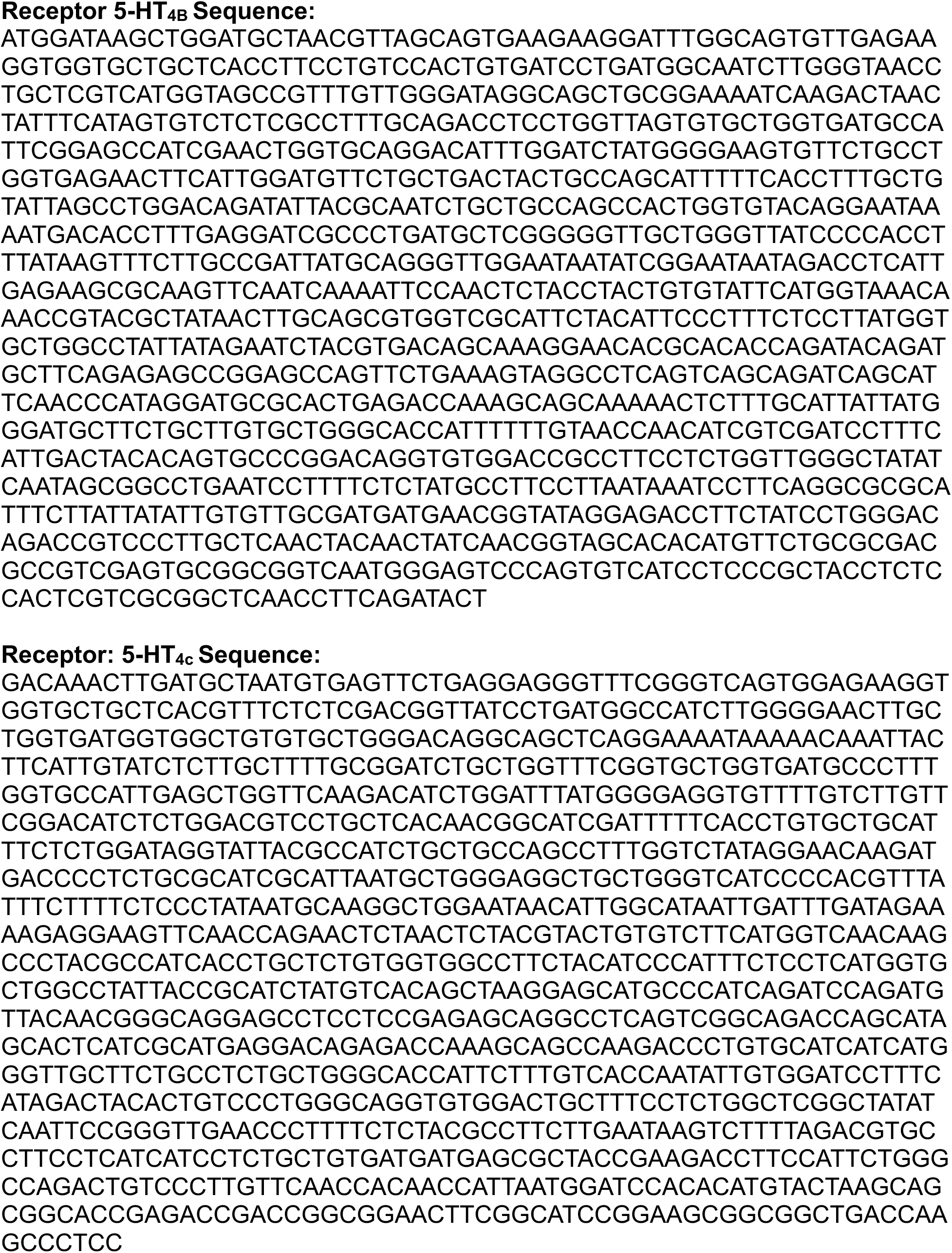
Isoform sequencing.

**Table S5.**
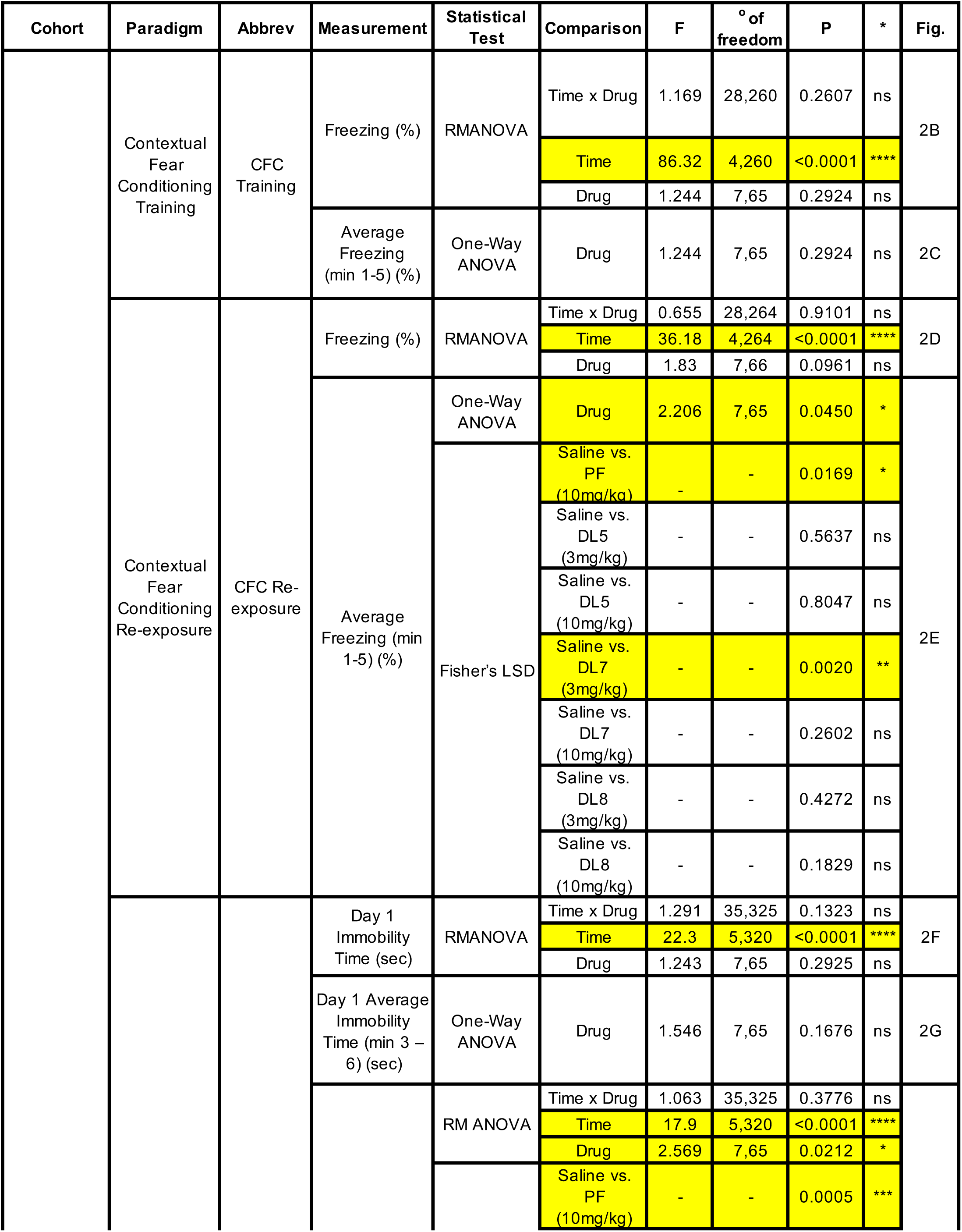
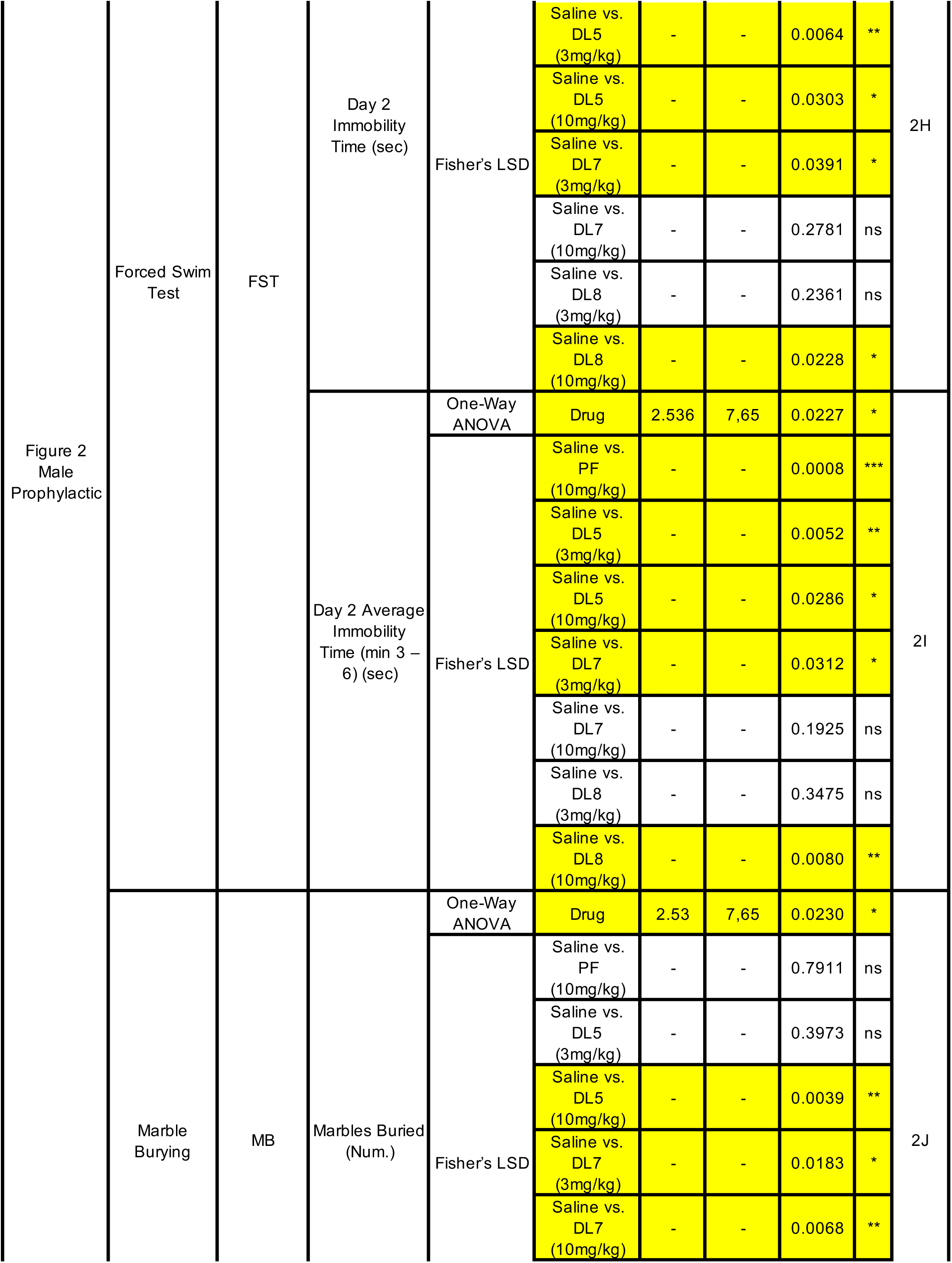
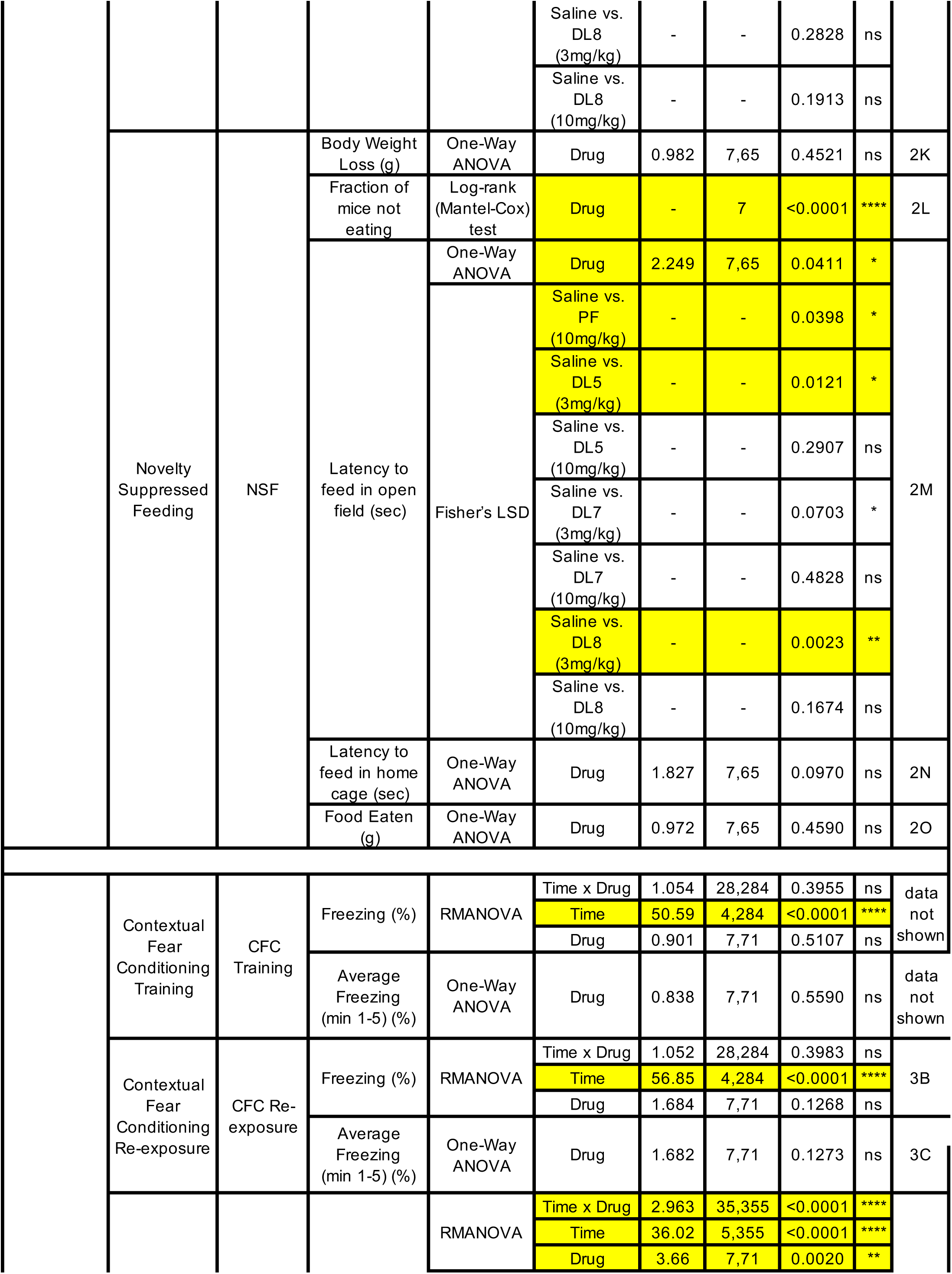
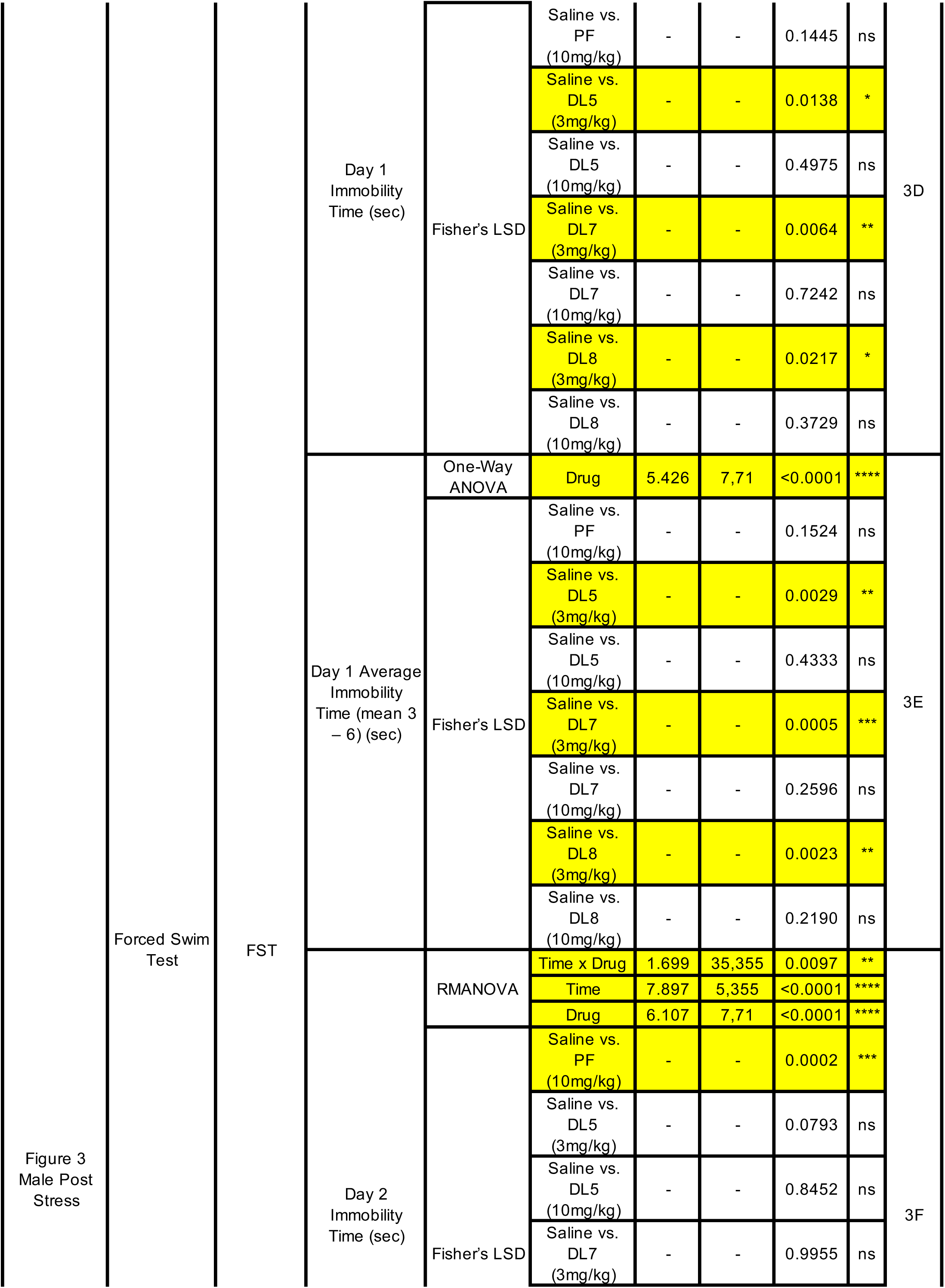
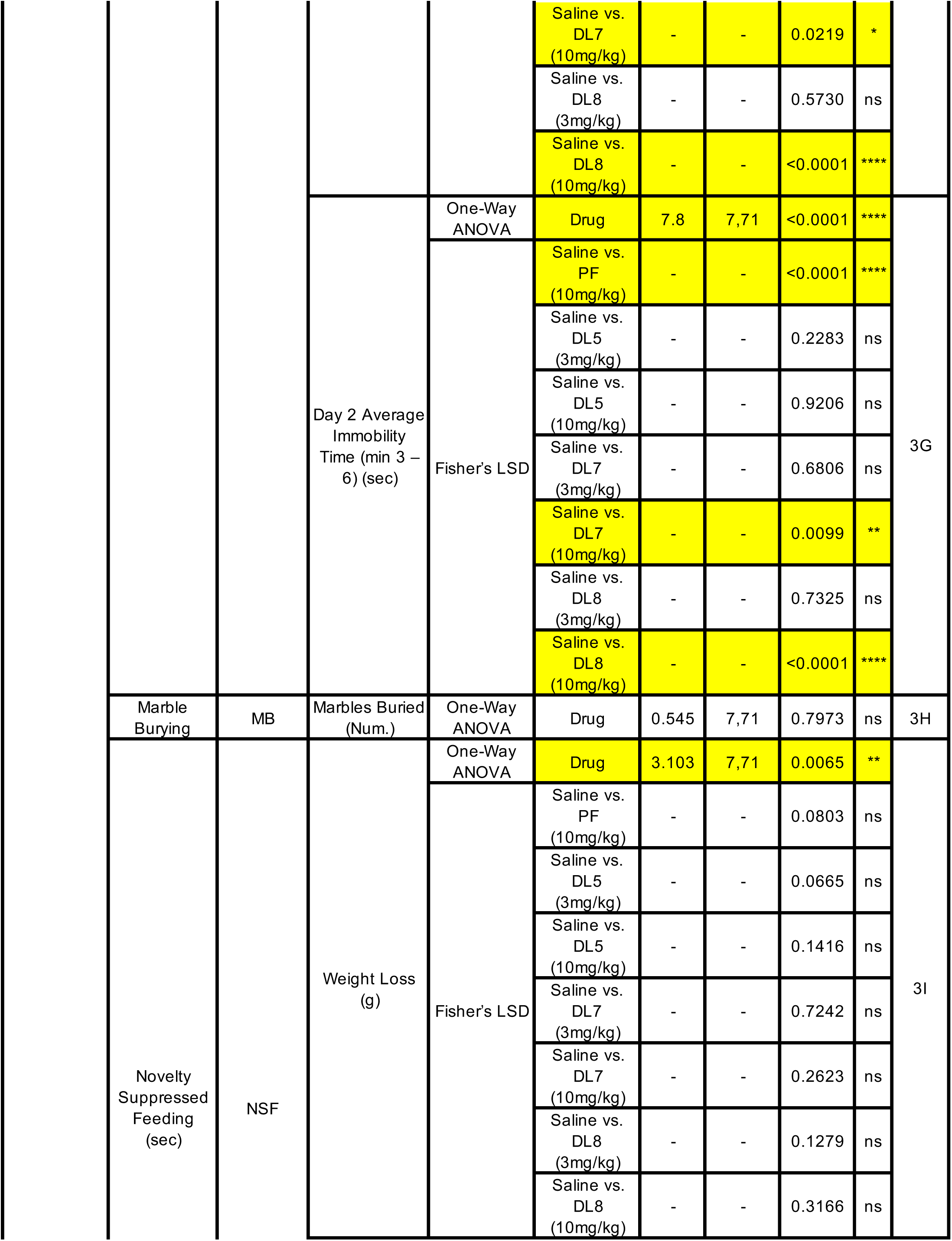
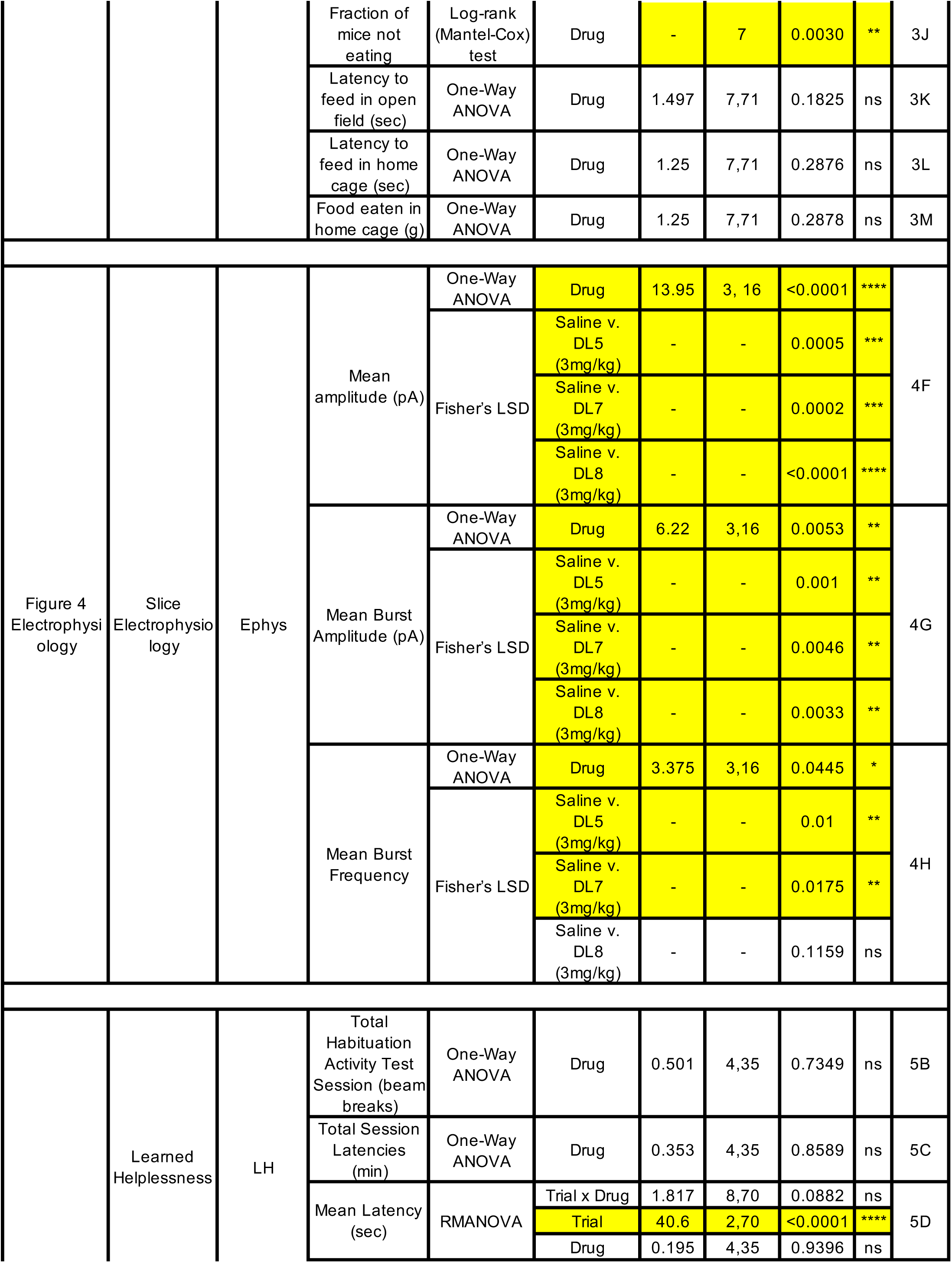
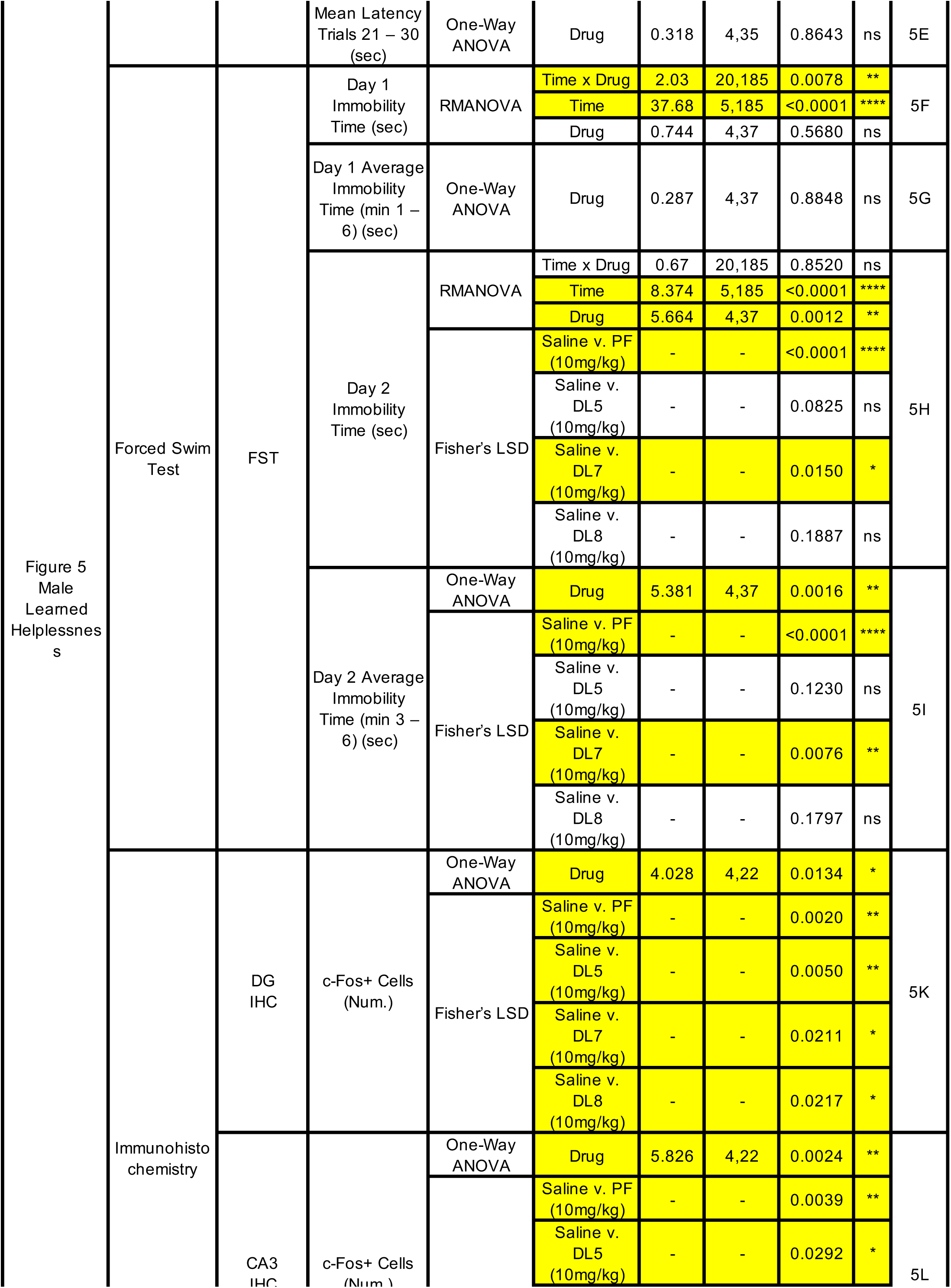
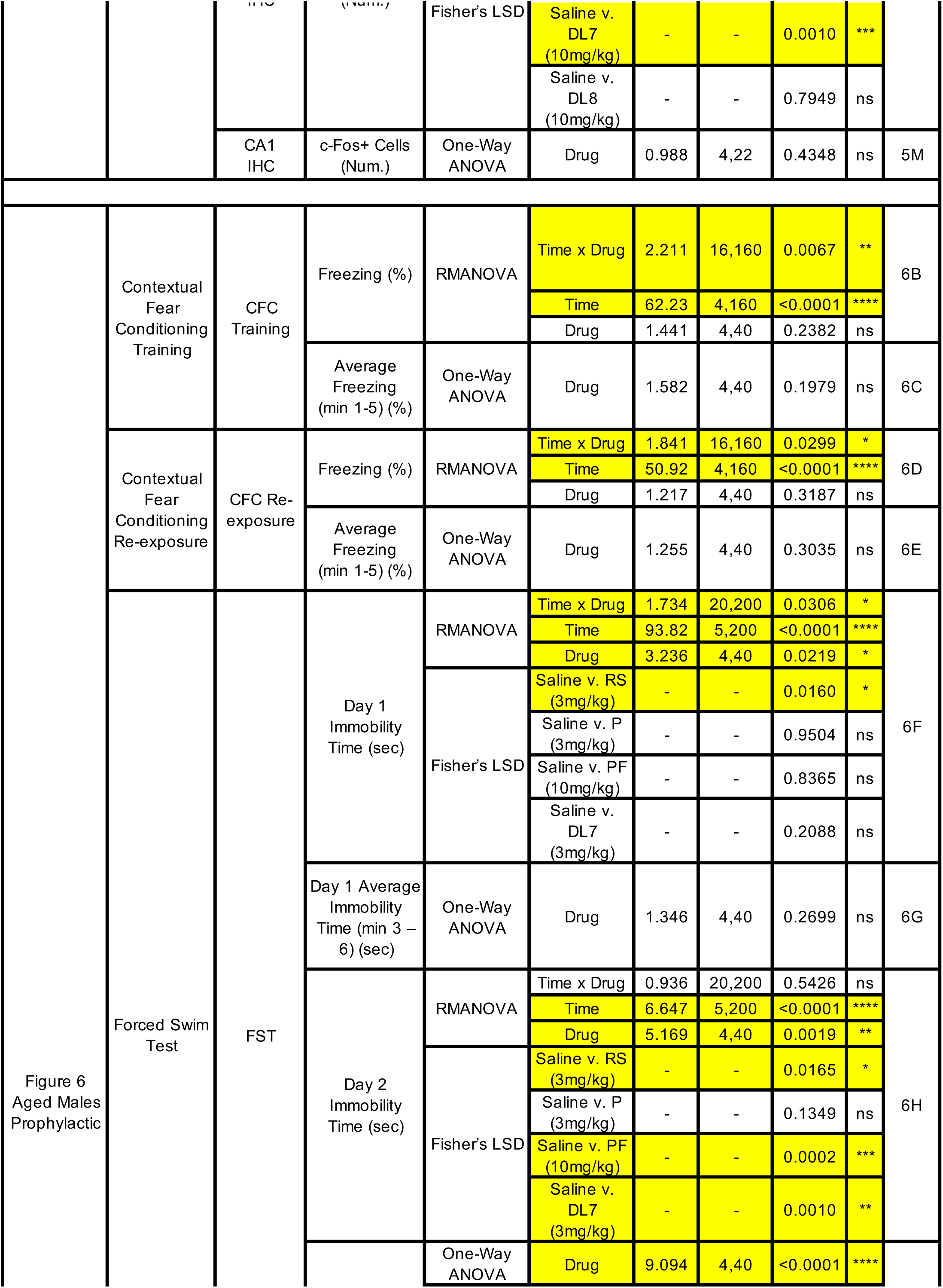
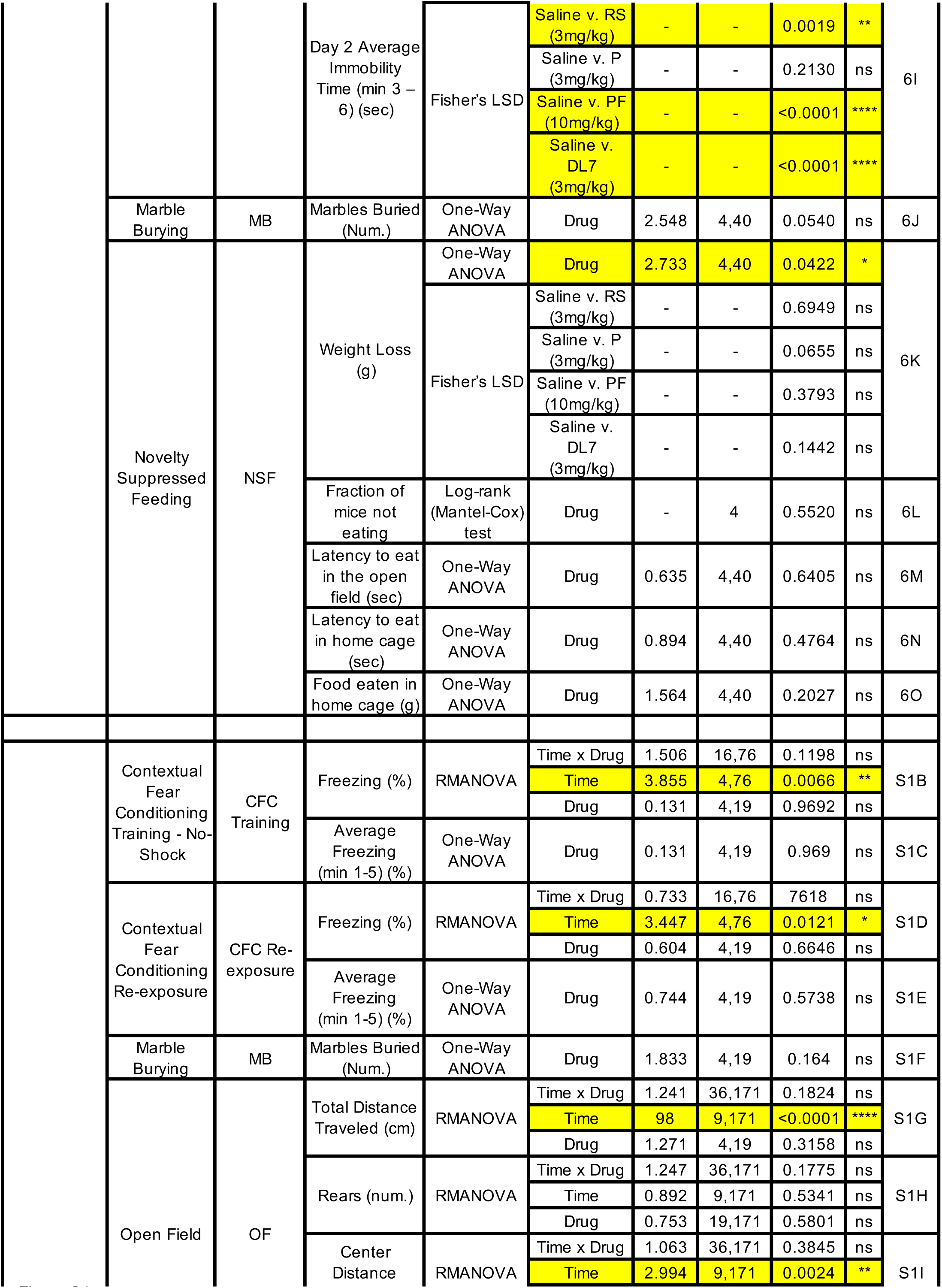
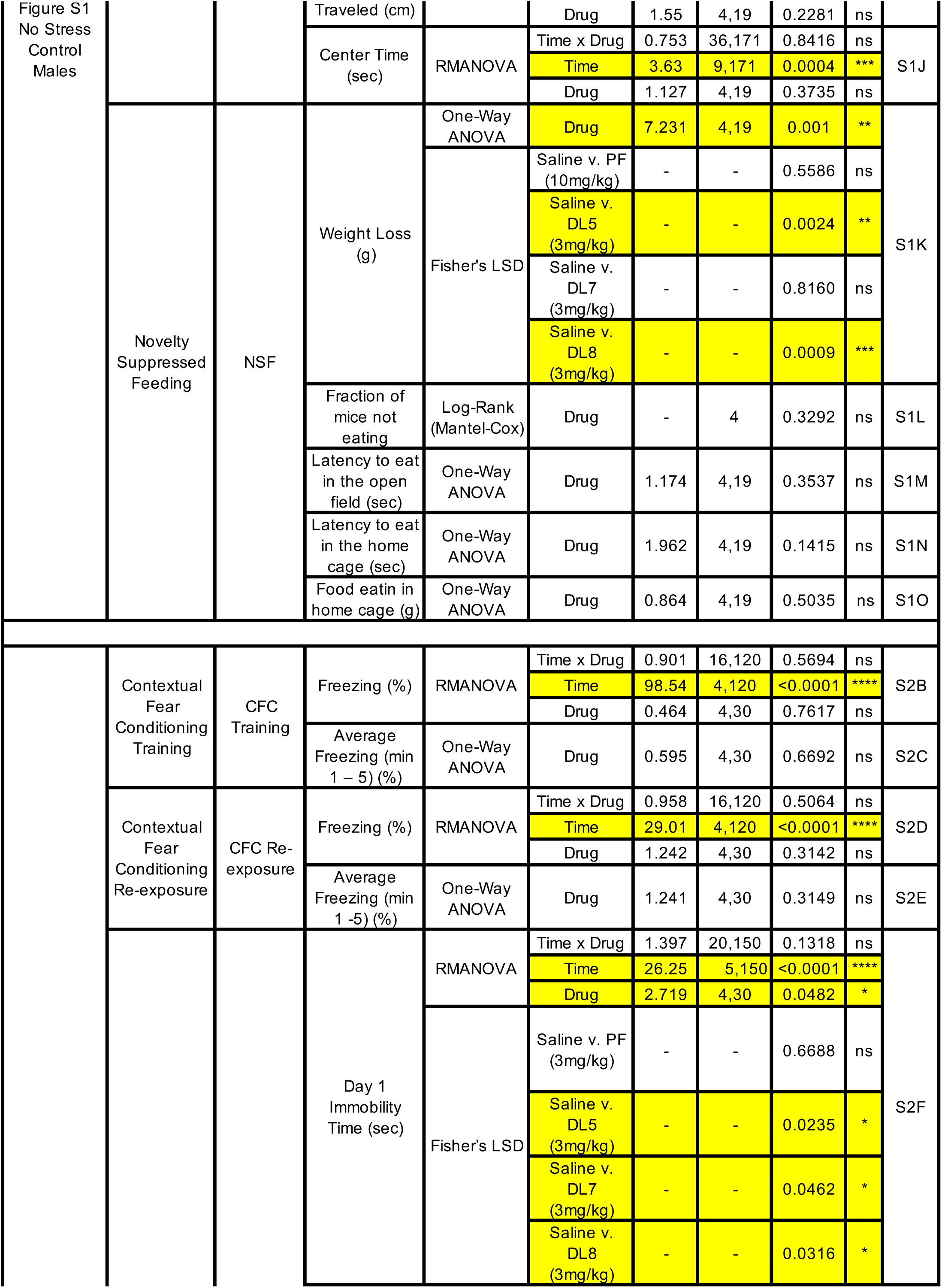
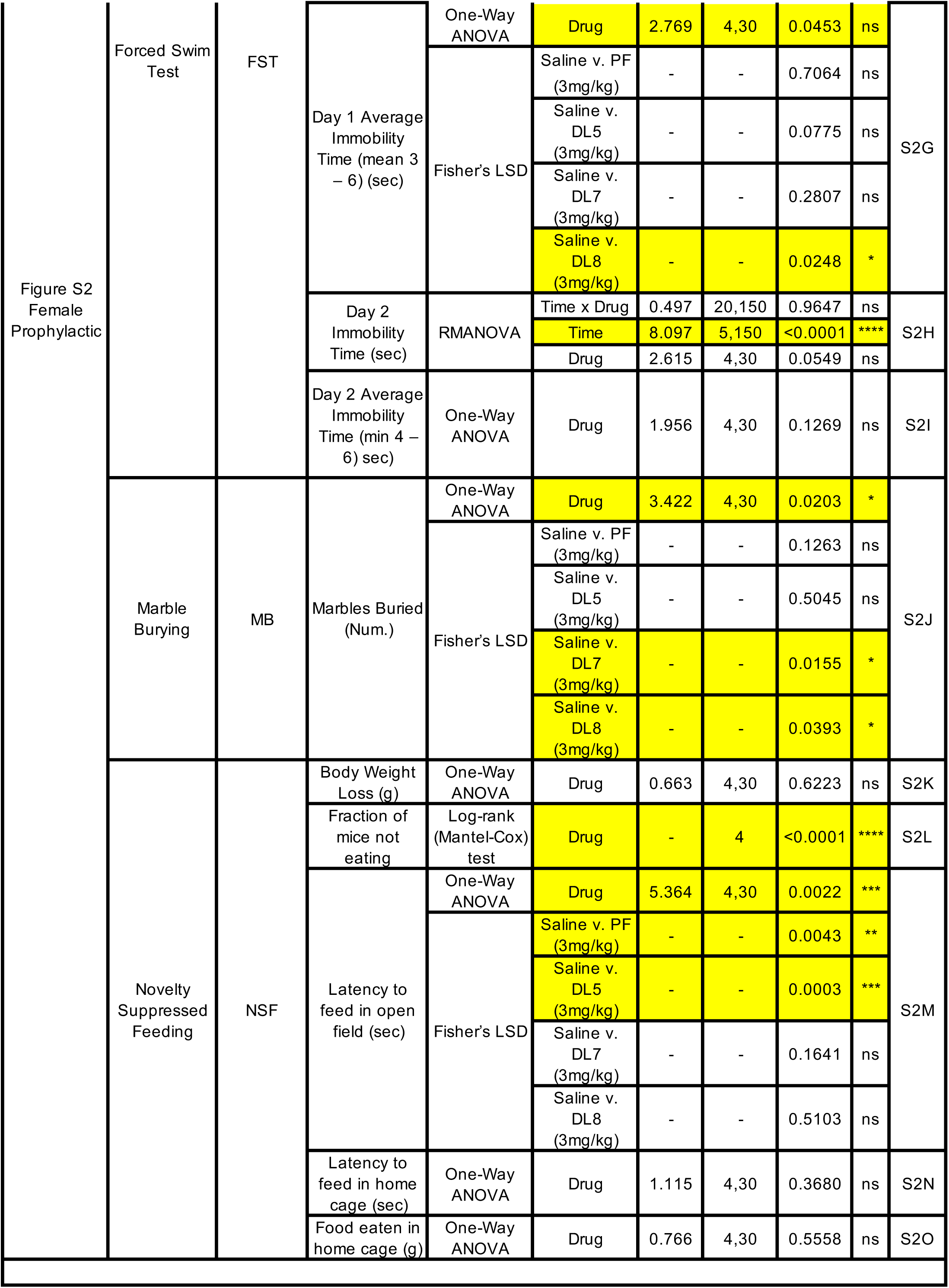
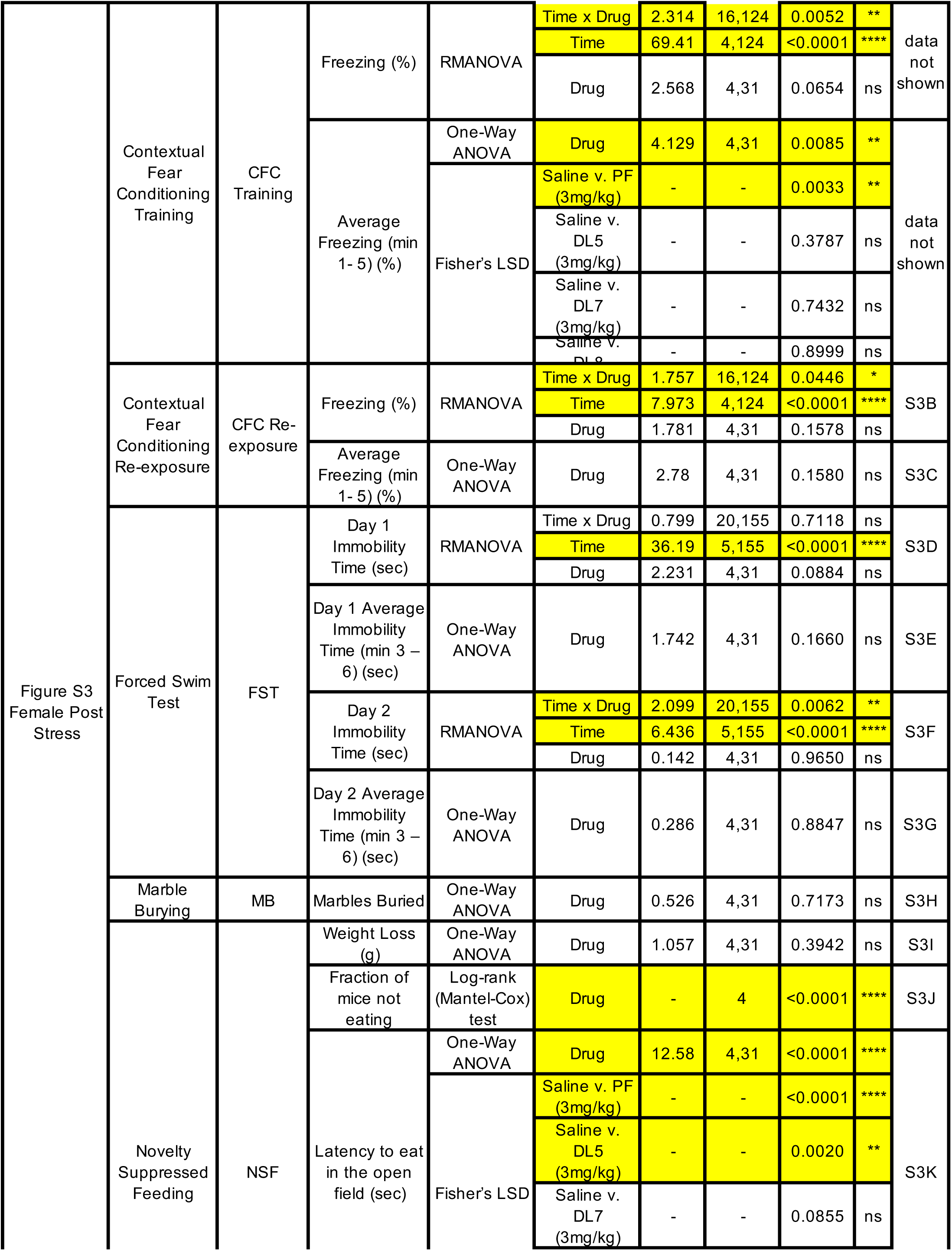
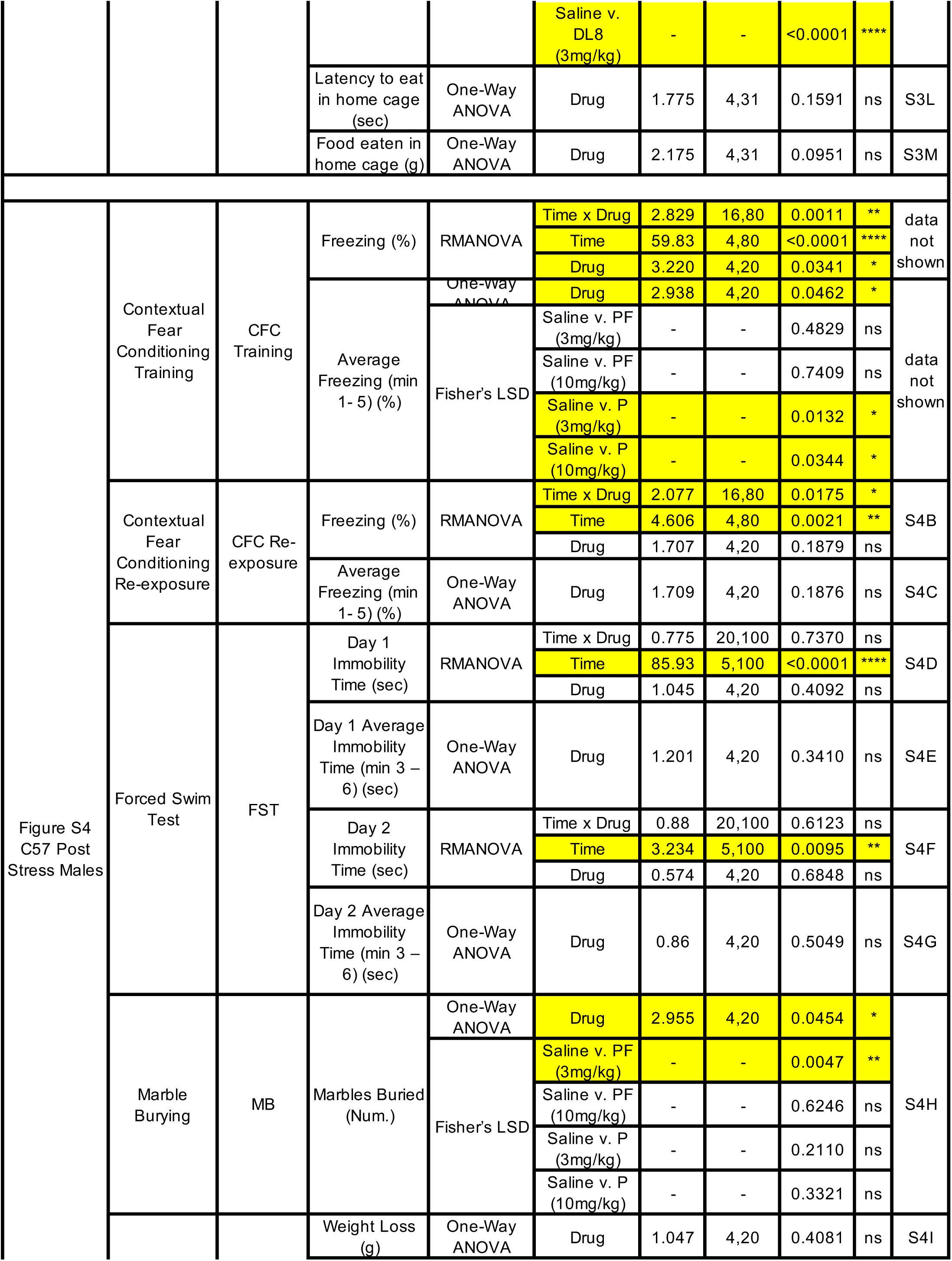
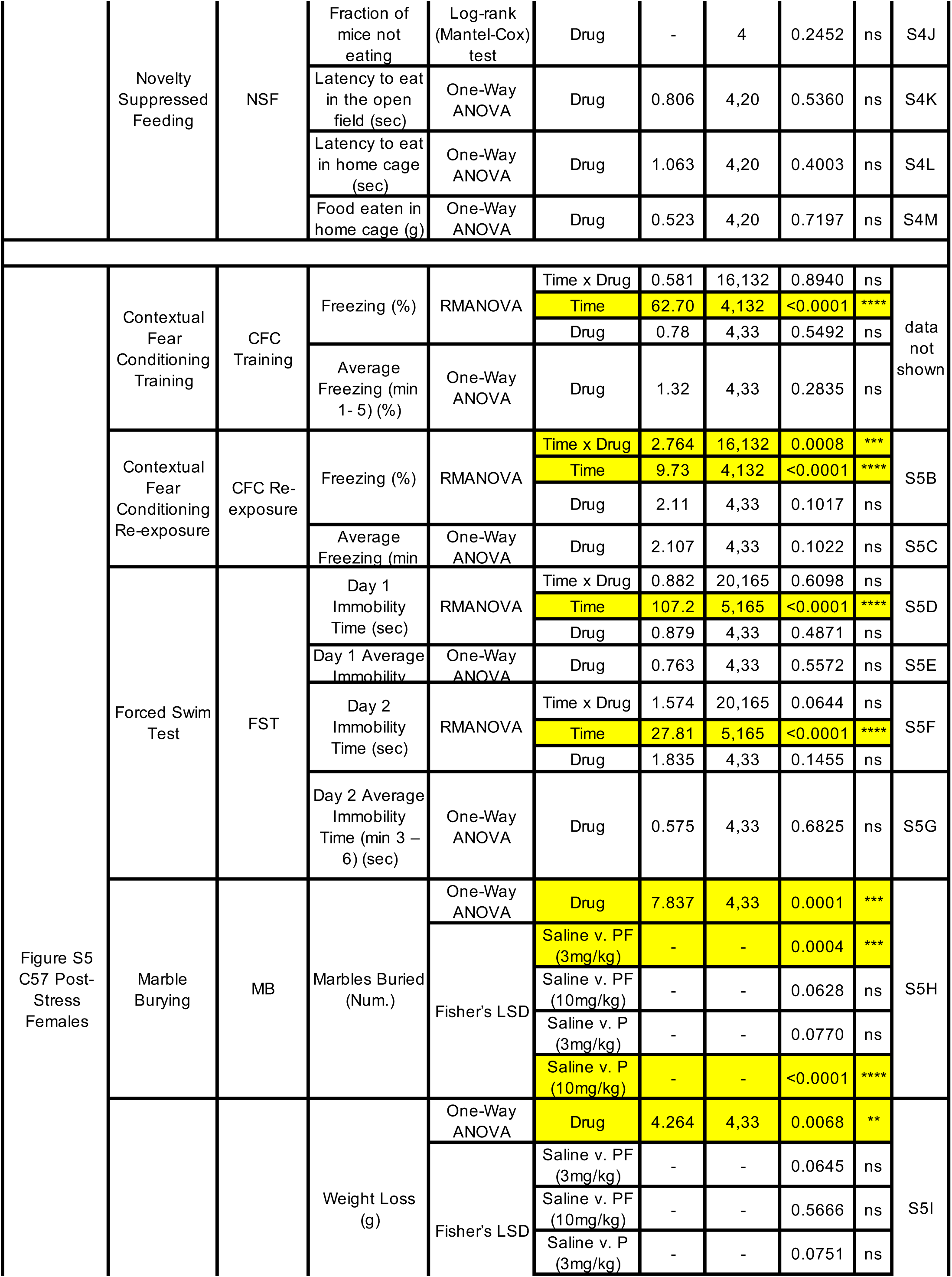
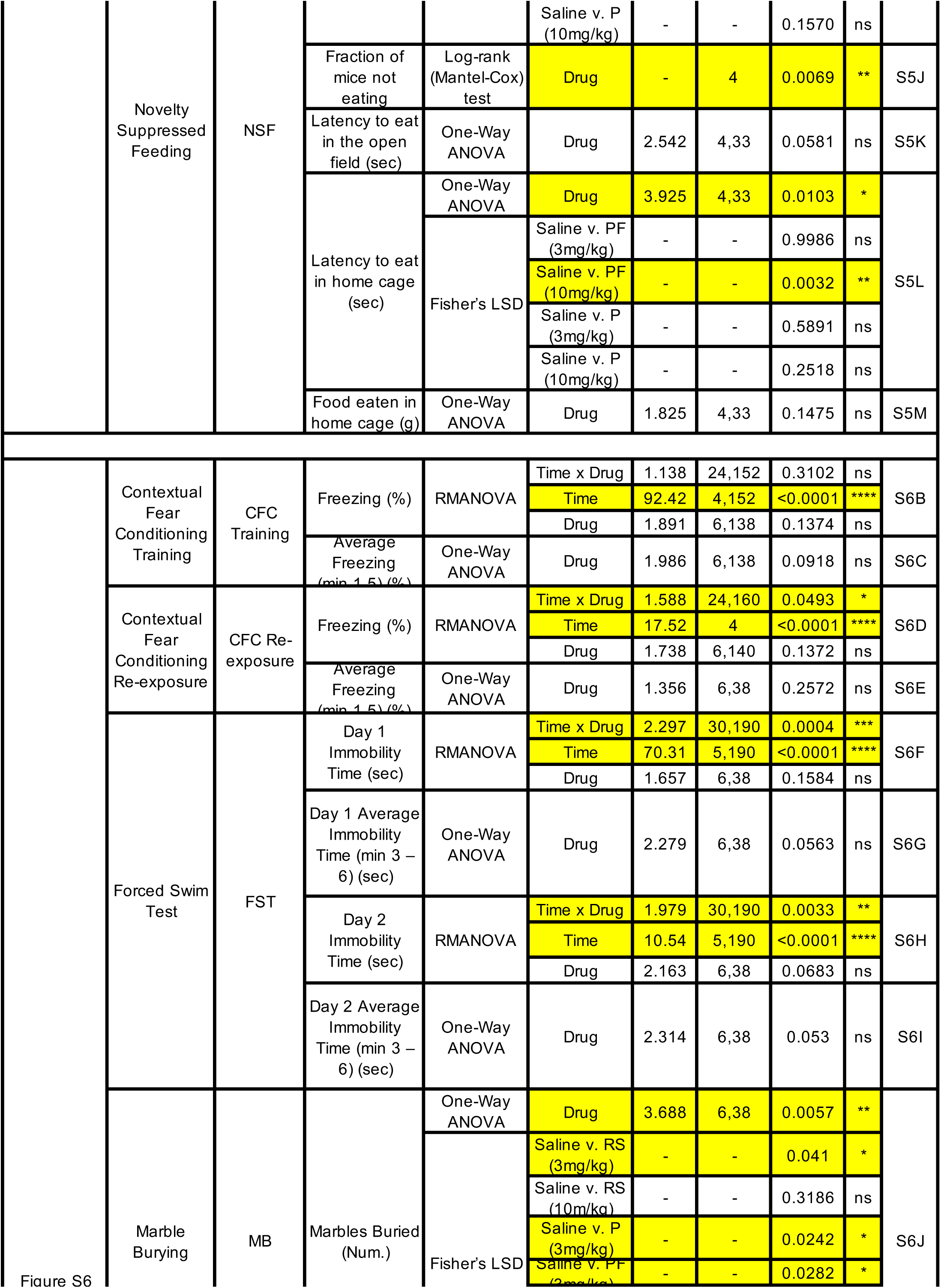
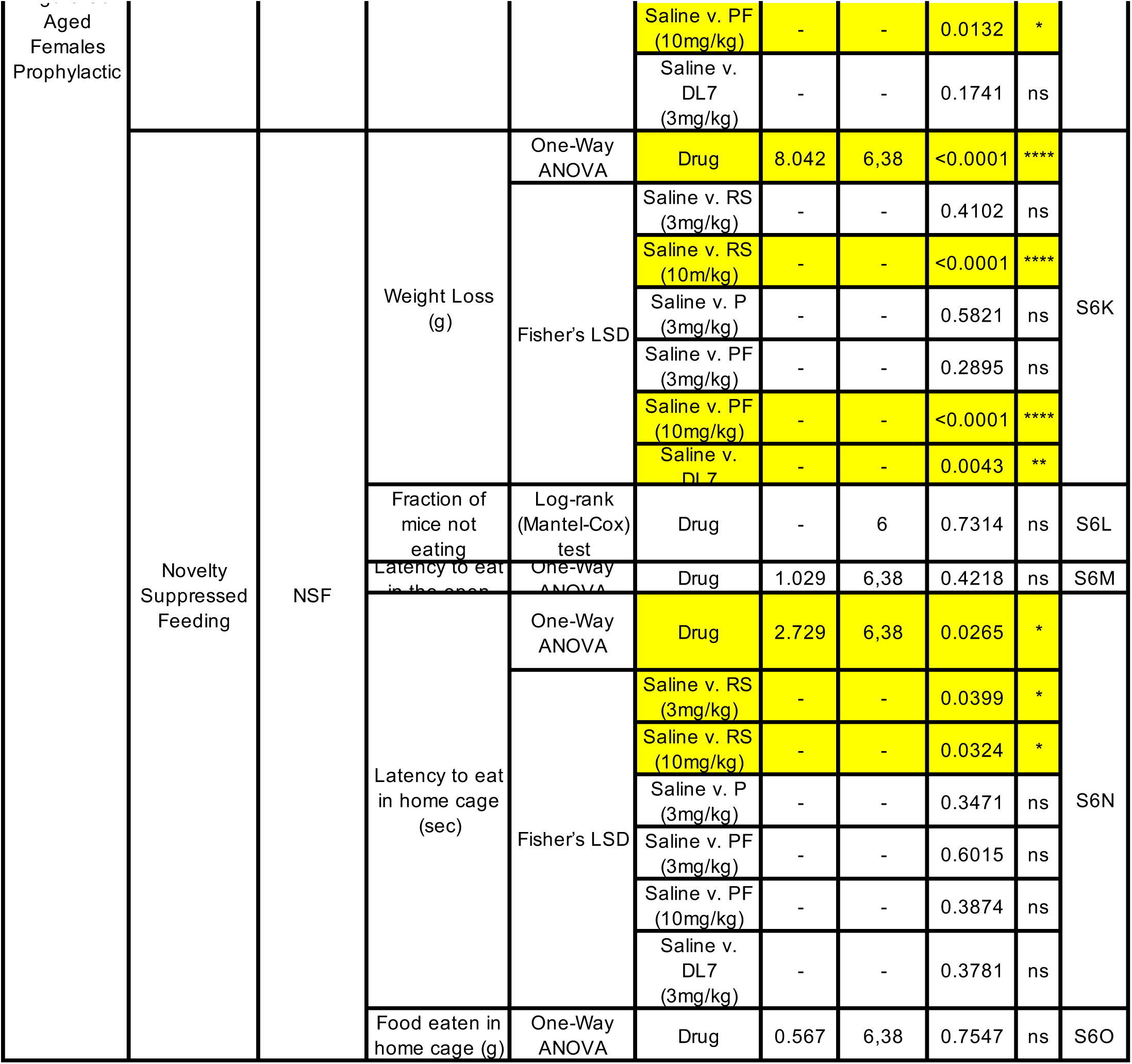
Statistical analysis summary for behavioral tests, electrophysiology, and c-Fos expression.

